# An Essential Role of the AMPK-Related Kinase SNRK in Regulating Drosophila Sleep

**DOI:** 10.64898/2026.04.29.721761

**Authors:** Weiwen Yang, Chaoyi Li, Jingyi Shi, Renbo Mao, Xinyue Xiong, Shichao Fan, Wenxia Zhang, Yi Rao

## Abstract

Protein kinases are important regulators of sleep in animals, but their loss-of-function mutations usually show moderate phenotypes. We report here that sucrose non-fermenting related kinase (SNRK), a member of the AMP-activated protein kinase-related kinase (ARK) family, plays an essential role in sleep regulation: *Snrk* gene knockout resulted in a profound loss of night-time sleep phenotype, much stronger than that of the well-known ARK member *Sik3*. This phenotype was completely rescued by neuronal SNRK expression. SNRK in adulthood was required for sleep. Catalytic and regulatory sites in SNRK were important for sleep. A chemoconnectomic (CCT) screen revealed SNRK functioning in cholinergic neurons whose activation is known to inhibit sleep. Sleep loss in SNRK mutants was rescued by pharmacological blockade of nicotinic acetylcholine receptors. Our findings have uncovered a fundamental role for SNRK in cholinergic neurons to regulate sleep.

## INTRODUCTION

Sleep is an important behavioral state in animals^1,2^. Research has been carried out to understand neural circuits^3–8^ and molecules^9–12^ involved in regulating sleep.

At the intracellular level, protein phosphorylation has been implicated in sleep regulation^13–22^. Several protein kinases have been found to regulate sleep: protein kinase A (PKA)^23,24,25^, the extracellular signal-regulated kinase (ERK)^18,26,27^, the adenosine monophosphate (AMP)-activated protein kinase (AMPK)^28–30^, calcium-calmodulin kinase (CaMK) II αandβ^31–33^, c-Jun N-terminal kinase (JNK)^34^, Anaplastic Lymphoma Kinase (ALK)^35^, and the liver kinase B (LKB1)^36^. A previous report of drastic sleep reduction in Camk2b knockout mice^32^ was later refuted by our work which supported a minor role for Camk2a but no involvement of Camk2b in basal sleep regulation^37^.

An unbiased forward-genetic screen led to the discovery of SIK3^15,38^, a member of the AMPK-related kinase (ARK) family. Specifically, gain-of-function (GOF) SIK3 mutations showed severe hypersomnia in mice^14,16,38–40^. Its upstream regulator, the kinase LKB1, was also found to be involved in regulating sleep^14,16,36^. However, loss-of-function (LOF) mutants of SIK3 in Drosophila and the mouse showed very moderate sleep phenotype^41–44^. No sleep phenotype was observed in LOF mutants of genes closely related to SIK3, such as SIK1 and SIK2^45^. These raised the question of the functional roles of other ARKs in sleep regulation.

*Drosophila melanogaster* provides a robust genetic model to address this question^46–49^. We conducted a reverse-genetic screen of the ARK family and found a strong reduction of nighttime sleep caused by LOF mutations of sucrose non- fermenting related kinase (SNRK)^50–54^. This phenotype depends on SNRK in neurons because it could be completely rescued by re-introduction of SNRK into neurons after it was knocked out from the entire body. Re-introduction to adult mutants rescued sleep and RNAi in adult flies reduced sleep, both supporting a role of SNRK functioning in adults, not in development. Mutations of either the catalytic sites or the regulatory sites in SNRK could reduce sleep. SNRK-knock-in flies allowed us to observe SNRK expression in neurons. To dissect neurons in which SNRK functions to regulate sleep, we used drivers of the chemoconnectome (CCT) which we created previously for functional studies of neurotransmission^55,56^. Sleep phenotype was found when SNRK RNAi was expressed in cholinergic neurons, whose activation was known to inhibit sleep^3,57–59^. Expression of SNRK in cholinergic neurons reversed sleep loss caused by SNRK mutations. Pharmacological inhibition of nicotinic acetylcholine receptors (nAChRs) also reversed severe insomnia in SNRK mutants. We have therefore discovered an essential role for SNRK functioning in cholinergic neurons to regulate sleep.

## RESULTS

### Profound Sleep Loss in SNRK LOF Mutants

To identify novel regulators of sleep in the ARK family (Figure 1A), we utilized homologous recombination to replace the endogenous locus of each kinase with a 3P3-red fluorescent protein (RFP) cassette to generated null alleles (*attP* lines) for the targeted ARKs (Figure 1B). Null alleles of *Ampk*, *Par-1* and *Nuak* (*CG43143*) resulted in lethality. Behavioral analysis of the LOF mutants for other ARKs revealed that, of all the viable ARK mutant flies, SNRK homozygote mutants (*Snrk-attP*) exhibited the most dramatic reduction in total sleep duration compared to genetic controls (*IsoCS*) (Figure 1C). SNRK mutant flies showed the most severe sleep reduction at nighttime.KP78β and SFF mutant flies showed moderate nighttime sleep phenotype. Daytime sleep reduction was most severe in Sff mutants (Figure 1C).

**Figure 1.**
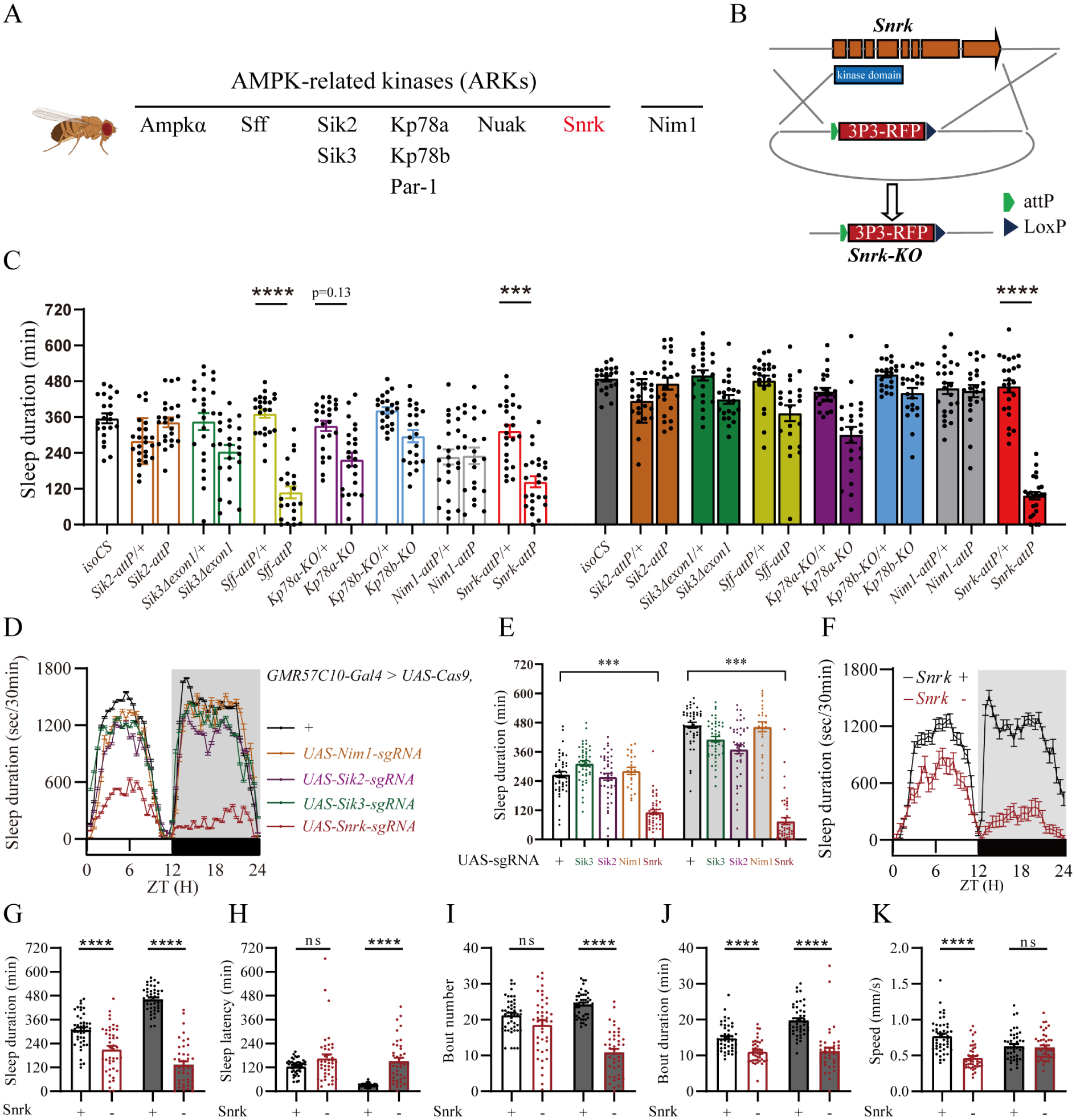
SNRK Regulation of Sleep in *Drosophila*. **(A)** ARK family members in *Drosophila*. **(B)** A schematic representation of the homologous recombination strategy used to generate the SNRK null mutant. The endogenous locus was replaced with a 3P3-RFP reporter cassette. **(C)** Behavioral screening of null or hypomorphic mutants of ARKs. Daytime (left panels) and night-time (right panels) sleep duration (min) of homozygous mutants and their respective genetic or heterozygous controls are shown. **(D)** Continuous 48-hour sleep profile following pan-neuronal somatic knockout of targeted ARK members (*Nim1*, *Sik2*, *Sik3*, *Snrk*). Knockout was induced by driving Cas9 expression pan-neuronally (*GMR57C10-Gal4 > UAS-Cas9*) alongside the respective *UAS-sgRNA* lines. **(E)** Quantification of daytime and night-time sleep duration (min) for the pan-neuronal somatic knockout flies and their transgenic controls. **(F)** A representative 24-hour sleep profile (Sleep duration sec/30min) of control (*Snrk* +) and SNRK null mutants (*Snrk* -) under 12 h light : 12 h dark cycles (ZT, Zeitgeber Time). **(G)** Quantification of total daytime and night-time sleep duration (min). **(H–K)** Detailed quantitative analysis of sleep architecture parameters during the day and night, including sleep latency (min) **(H)**, sleep bout number **(I)**, and sleep bout duration (min) **(J)**. **(K)** Quantification of waking speed (mm/s) to assess locomotor activity during wakefulness. ns, not significant; *p < 0.05; **p <0.01; ***p <0.001 and ****p <0.0001; mean ± standard error of the mean (mean ± SEM). Kruskal-Wallis test **(C, E)**; Mann-Whitley test **(G, H, I, J, K).**

To bypass developmental lethality^41^ and assess LOF phenotypes in the nervous system, we used an *in vivo* pan-neuronal CRISPR/Cas9 somatic knockout strategy ^55^ (Figure 1 D and E). We targeted NIM1, SIK2, SIK3 and SNRK (Figure S1A). The sleep-loss phenotype in observed in pan-neuronal disruption of SNRK was stronger than those of NIM1, SIK2 or SIK3 (Figure 1 D and E, Figure S1 B and C).

While daytime sleep was moderately decreased, night-time sleep was nearly abolished in SNRK null mutants (Figure 1F and G). This was due to defects in sleep initiation and maintenance. SNRK deficiency increased latencies of nighttime but not daytime sleep (Figure 1H), reduced bout numbers of nighttime but not daytime sleep (Figure 1I), and reduced bout durations of nighttime as well as daytime sleep (Figure 1J). Walking speed analysis showed that SNRK mutants were not hyperactive: their locomotion speed was lower than the wild type (WT) during the day and indistinguishable from the WT at night (Figure 1K).

To examine whether sleep loss was secondary to defective circadian rhythmicity, we analyzed the locomotor activities of SNRK mutants under constant darkness (DD). Both single-fly actograms and population activity profiles showed no circadian defects in SNRK mutants (Figure S2 A, B and C). Continuous monitoring under free-running conditions confirmed that the sleep reduction persisted independently of light cues (Figure S2 D and E). SNRK mutants exhibited a normal free-running circadian period statistically indistinguishable from wt flies (Figure S2F). Thus, it is unlikely that the sleep phenotype in SNRK mutants was due to circadian defects.

### Rescue of SNRK Sleep Deficit by Neuronal Expression of the WT SNRK but not a Kinase Dead SNRK Mutant

To exclude possible off-target effects of gene targeting and address whether SNRK in the nervous system was sufficient in sleep regulation, we performed genetic rescue experiments using the pan-neuronal driver GMR57C10-Gal4 to restore WT SNRK expression specifically in neurons in the SNRK null mutant background (V: *Snrk^-/-^, GMR57C10-Gal4>UAS-Snrk*).

Pan-neuronal expression of WT SNRK completely rescued the sleep reduction in SNRK null mutants. Total sleep in the 24-hour profile, daytime sleep duration and night-time sleep duration were all rescued to the level of the WT when SNRK was re-introduced into neurons in SNRK null flies (Figure 2 A and B). Control flies carrying either the Gal4 driver alone (II: *Snrk^-/-^, GMR57C10-Gal4/+* in Figure 2) or the UAS responder alone (III: *Snrk^-/-^, UAS-Snrk/+* in Figure 2) in the mutant background had severe sleep phenotypes (Figure 2 A and B). Detailed analysis showed that pan-neuronal re-introduction of SNRK completely normalized the sleep bout number, bout duration, and sleep latency (Figure 2 C, D and E).

**Figure 2.**
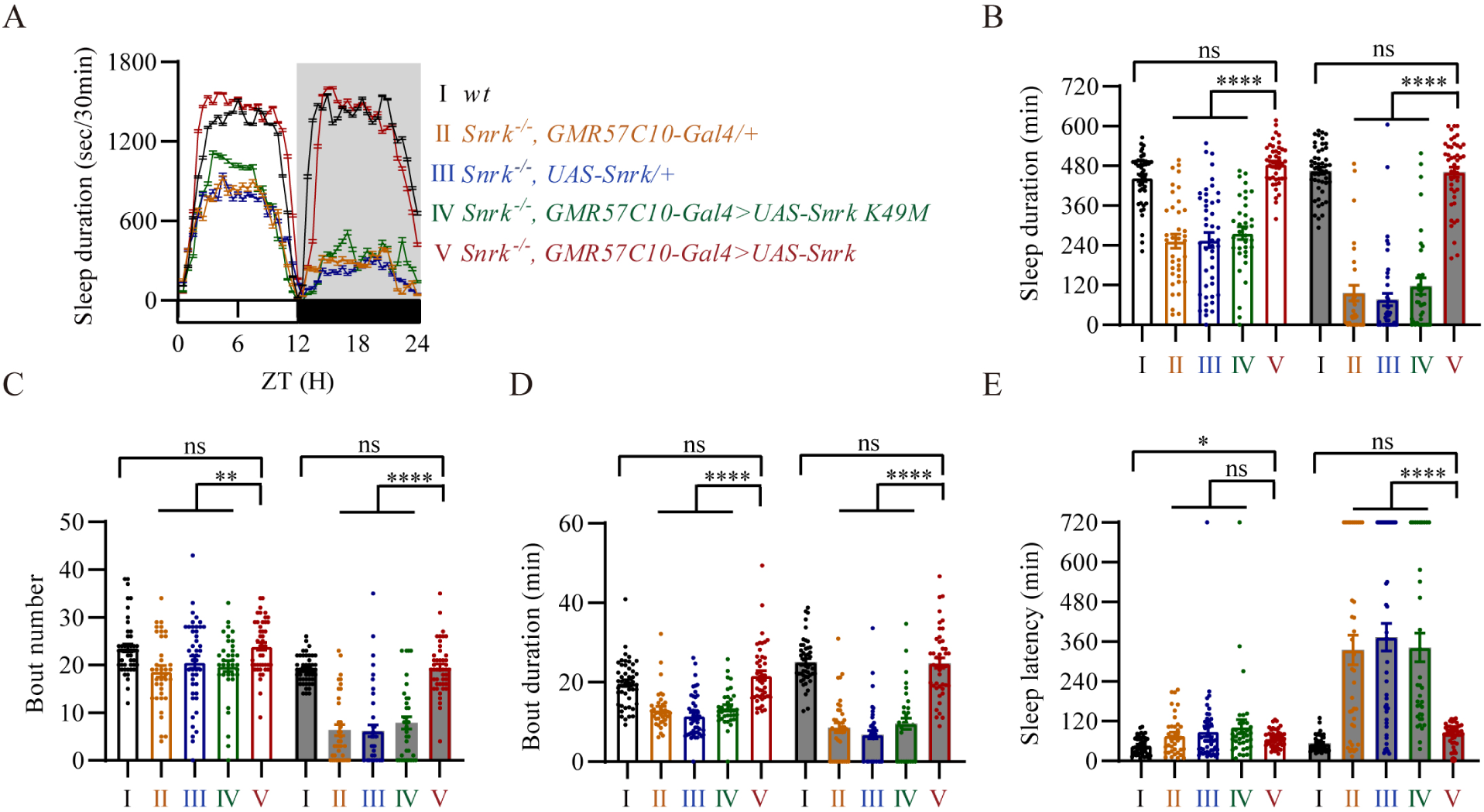
Sleep Rescue by Pan-neuronal Expression of SNRK in a Kinase-Dependent Manner. **(A)** A 24-hour sleep profile. The experimental and control groups are indicated as follows: wild-type control (I: *wt*), Gal4 driver control in the mutant background (II: *Snrk^-/-^, GMR57C10-Gal4/+*), UAS responder control in the mutant background (III: *Snrk^-/-^, UAS-Snrk/+*), kinase-dead rescue (IV: *Snrk^-/-^,GMR57C10-Gal4>UAS-Snrk-K49M*), and wild-type rescue(V: *Snrk^-/-^, GMR57C10-Gal4>UAS-Snrk*). **(B)** Quantification of daytime (left panels) and night-time (right panels) sleep duration (min). **(C–E)** Detailed quantitative analysis of sleep architecture parameters during the day and night, including sleep bout number **(C)**, sleep bout duration (min) **(D)**, and sleep latency (min) **(E)**. ns, not significant; *p < 0.05; **p <0.01; ***p <0.001 and ****p <0.0001; mean ± standard error of the mean (mean ± SEM). Kruskal-Wallis test **(B, C, D, E).**

In these experiments, we also investigated whether the kinase activity of SNRK was required for its sleep regulating function. We introduced a kinase-dead form of SNRK carrying a mutation (lysine to methionine) in its catalytic domain (at amino acid residue 49) (*UAS-Snrk K49M*). Pan-neuronal expression of the kinase-dead variant (IV: *Snrk^-/-^, GMR57C10-Gal4>UAS-Snrk K49M* in Figure 2) failed to rescue sleep loss in SNRK mutants (Figure 2A). Total sleep duration in 24 hours, sleep bout number, bout duration, and sleep latency could not be rescued by the kinase-dead SNRK mutant (Figure 2 B, C, D and E).

These results demonstrate that SNRK functions within the nervous system to regulate sleep, and that its enzymatic kinase activity is indispensable for sleep regulation.

### Sleep Regulating Function of SNRK in the Adult Nervous System

To rule out the possibility that sleep loss in SNRK mutants was a secondary consequence of developmental abnormalities, we investigated the temporal requirement of SNRK with both rescue and knockdown types of manipulations during adulthood.

We first used the inducible GeneSwitch (GS) system^60–62^. The *elavGS* driver was used to express a SNRK transgene in the pan-neuronal population of adult SNRK null mutants with the inducer RU486 applied to adult flies. Adult-specific pan-neuronal expression of wt SNRK restored the daily sleep profile and total sleep duration (Figure 3 A and B). Adult induction of wt SNRK rescued sleep bout duration (Figure S3A) and sleep bout number (Figure S3C). By contrast, in uninduced flies (without RU486 treatment), all experimental and transgenic control groups showed defective sleep (Figure S3C). Furthermore, adult-specific expression of the kinase-dead form (*UAS-Snrk K49M*) failed to rescue sleep defects (Figure 3 A and B). These data support that SNRK regulates sleep in adulthood, in a kinase-dependent manner.

**Figure 3.**
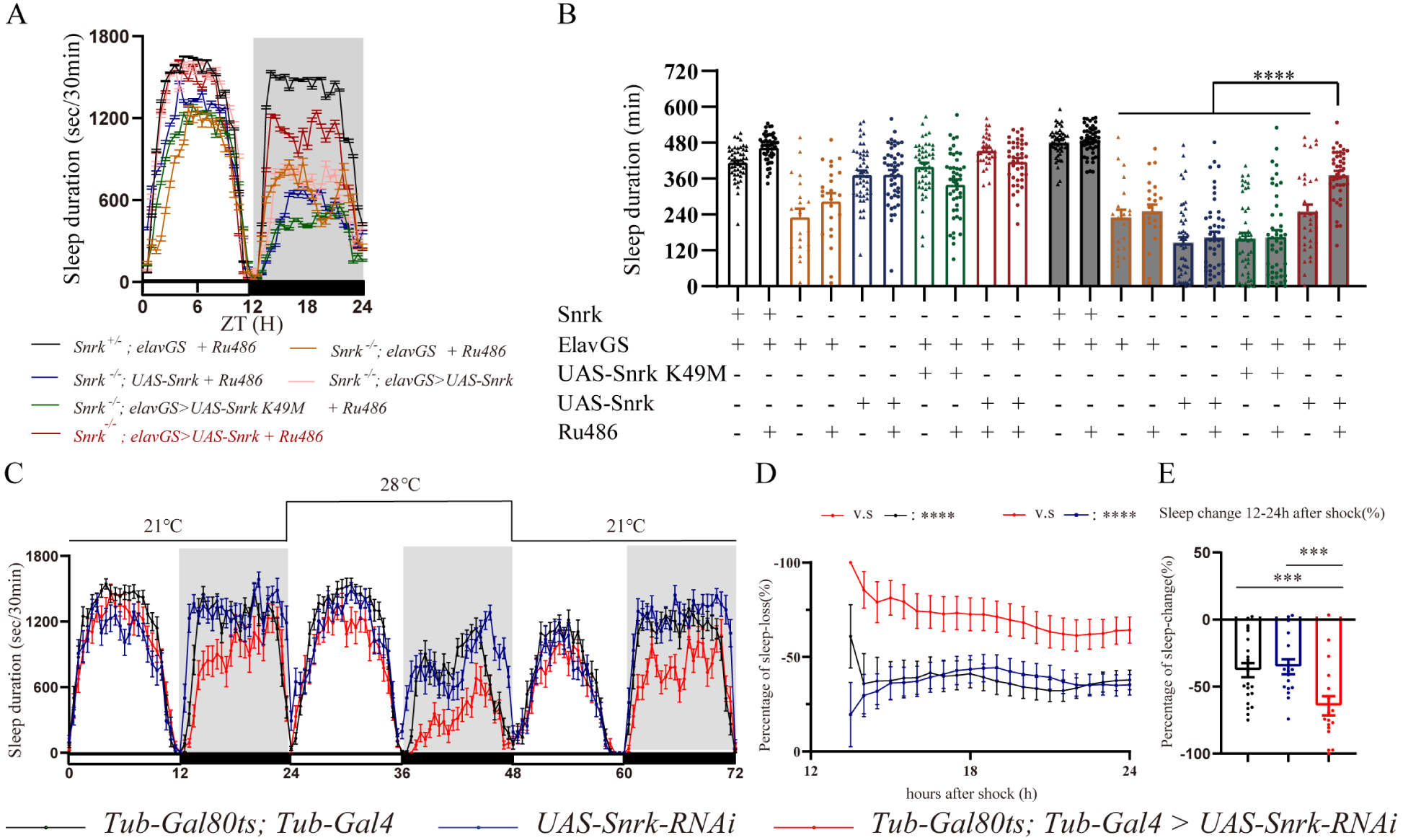
Requirement of Adult SNRK in Sleep Regulation. **(A)** A 24-hour sleep profile of adult flies utilizing the RU486-inducible GeneSwitch system (*elavGS*). Profiles are shown for SNRK null mutants expressing either wild-type SNRK or the kinase-dead variant (*UAS-Snrk K49M*) pan-neuronally, alongside corresponding controls, upon feeding with the RU486 inducer. **(B)** Quantification of total sleep duration (min) for the indicated genotypes in the presence (+) or absence (-) of RU486. **(C)** Continuous sleep profile (Sleep duration sec/30min) over a 72-hour period utilizing the temperature-sensitive TARGET system (*tub-Gal80ts; tub-Gal4*). Flies were initially maintained at the permissive temperature (21 °C), shifted to the restrictive temperature (28 °C) to induce SNRK RNAi knockdown, and subsequently returned to 21 °C. **(D)** Hourly tracking of the percentage of sleep loss (%) during the 12-hour restrictive temperature window (hours 14 to 24 after the temperature shock) for the SNRK knockdown group versus genetic controls. **(E)** Quantification of the overall sleep change (%) during the 12-24 hour period following the shift to 28°C, normalized to baseline sleep levels. ns, not significant; *p < 0.05; **p <0.01; ***p <0.001 and ****p <0.0001; mean ± standard error of the mean (mean ± SEM). Friedman test (**D**); Kruskal-Wallis test **(B, E).**

We then knocked down SNRK globally and specifically in adult flies using the temperature-sensitive Temporal and Regional Gene Expression Targeting (TARGET) system^63,64^ (Figure 3C). Flies expressing *tub-Gal80ts*; *tub-Gal4* and *UAS- SNRK RNAi* were similar to the wt in their sleep profiles when raised at the permissive temperature (21°C). However, shifting the environment to 28°C, which inactivated the Gal80ts repressor and allowed acute RNAi-mediated SNRK knockdown, significantly reduced sleep. Quantification revealed a significant percentage of sleep loss specifically in the RNAi-induced flies during the time following the temperature shock (Figure 3 D and E).

Thus, results from temporally controlled, bidirectional rescue and knockdown experiments confirmed adult requirement of SNRK in sleep regulation.

### SNRK Sequences Functionally Important for Sleep Regulation

The SNRK protein is conserved across flies, mice and humans (Figure S5A). To test the functional conservation *in vivo*, we introduced the human SNRK (hSNRK) into Drosophila. Pan-neuronal expression of the human SNRK gene (*UAS-hSNRK*) in the Drosophila SNRK null mutant background rescued the sleep defects. hSNRK expression restored the 24-hour sleep profile (Figure S4B), total sleep duration (Figure S4A), sleep bout number (Figure S4C) and sleep bout duration (Figure S4D).

Within the catalytic domain SNRK, there was a specific threonine residue at position 177 (T177) in Drosophila, corresponding to a putative regulatory site which was subject to phosphorylation ^65^. To investigate the physiological significance of T177 in sleep regulation, we used a PhiC31 integrase-mediated cassette exchange strategy. We replaced the endogenous SNRK locus with engineered point mutations, generating stable knock-in fly lines (Figure 4A): a wt sequence, a phosphomimetic mutation of glutamate replace of threonine (T177E), a phospho-resistant mutation of alanine replacement (T177A), and the kinase-dead mutation (K49M).

**Figure 4.**
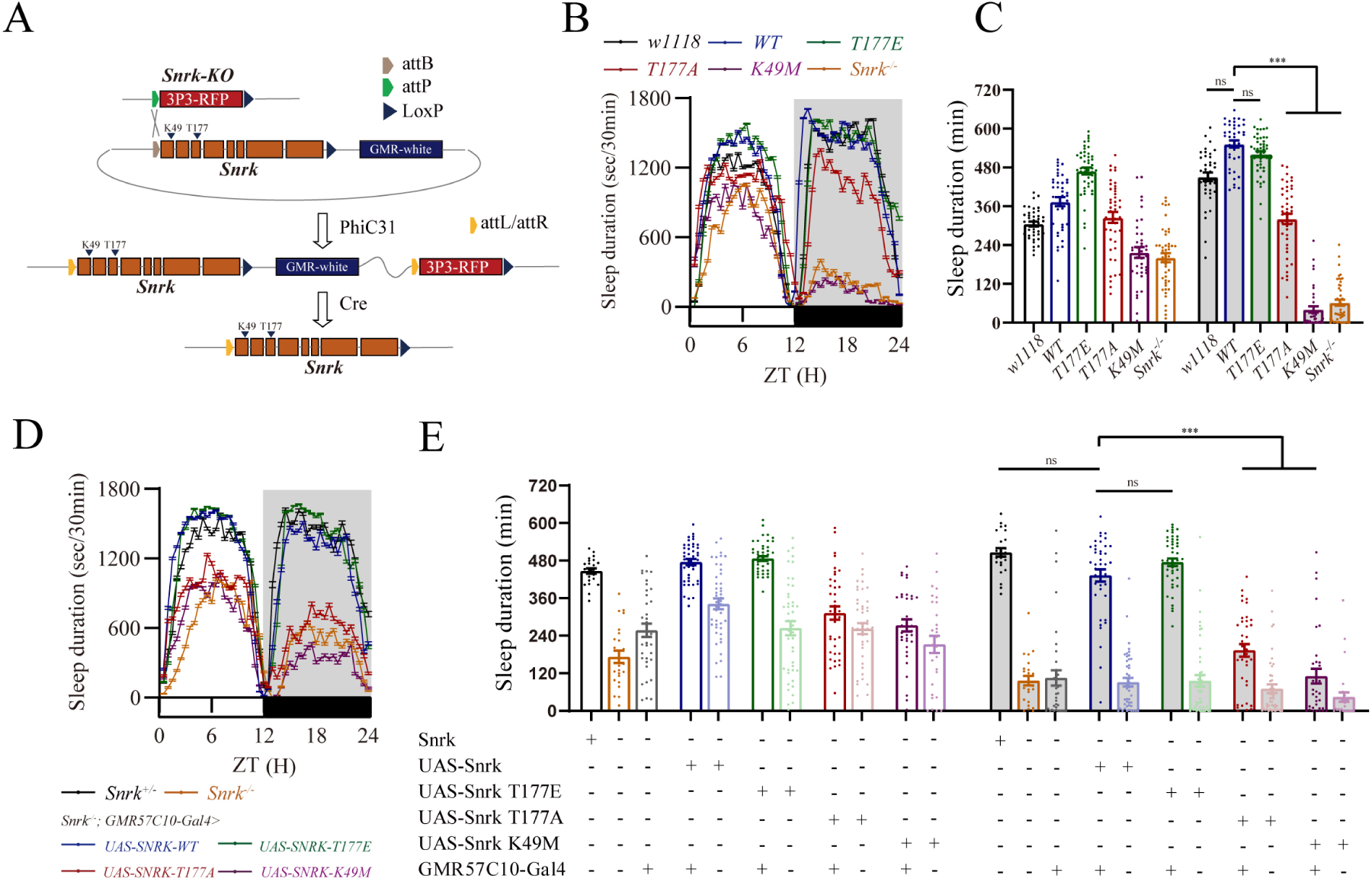
Requirement of Kinase SNRK Activity for Sleep Regulation. **(A)** A schematic representation of the PhiC31 integrase-mediated cassette exchange strategy used to generate endogenous *Snrk* knock-in lines. The 3P3-RFP reporter in the *Snrk-KO* locus was replaced with engineered alleles, including wild-type (*WT*), phosphomimetic (*T177E*), phospho-null (*T177A*), and kinase-dead (*K49M*) variants. **(B)** Continuous 48-hour sleep profile of the endogenous *Snrk* knock-in lines (*WT*, *T177E*, *T177A*, *K49M*). **(C)** Quantification of daytime and night-time sleep duration (min) for the indicated endogenous knock-in variants, alongside *w1118* (wild-type control) and complete SNRK null mutants. **(D)** Continuous 48-hour sleep profile of transgenic rescue flies expressing wild-type SNRK (*UAS-Snrk*), phosphomimetic (*UAS-Snrk T177E*), phospho-null (*UAS-Snrk T177A*), or kinase-dead (*UAS-Snrk K49M*) variants pan-neuronally (*GMR57C10-Gal4*) in the SNRK null mutant background. **(E)** Quantification of night-time sleep duration (min) for the pan-neuronal transgenic rescue groups described in (D) and their corresponding genetic controls. ns, not significant; *p < 0.05; **p <0.01; ***p <0.001 and ****p <0.0001; mean ± standard error of the mean (mean ± SEM). Kruskal-Wallis test **(C, E).**

Behavioral analysis of these knock-in lines revealed that flies carrying the wt SNRK or the phosphomimetic T177E allele exhibited normal 24-hour sleep profiles and durations (Figure 4 B and C). Both sleep bout number (Figure S5B) and sleep bout duration (Figure S5C) were statistically indistinguishable from wt controls. By contrast, the phospho-resistant T177A mutation significantly reduced night-time sleep, accompanied by a reduction in sleep bout duration (Figure S5C). Furthermore, the kinase-dead K49M mutant displayed a drastically loss of night-time sleep driven with severe fragmentation and reduced bout duration (Figure S5 B and C), phenocopying the sleep defects observed in the SNRK null mutants.

We further confirmed these specifically in the nervous system via transgenic rescue experiments using the pan-neuronal *GMR57C10-Gal4* driver in the SNRK null background (Figure 4 D and E, Figure S5D). Consistent with the knock-in results, pan-neuronal expression of the wt SNRK or the T177E variant completely rescued total sleep loss, normalizing both sleep bout number (Figure S5E) and bout duration (Figure S5F). The T177A variant was significantly impaired in its rescue capacity, resulting in shortened bout duration (Figure S5 E and F). The kinase-dead K49M variant completely failed to rescue the sleep phenotype (Figure S5 E and F).

### Expression and Function of SNRK in the Nervous System

To map the endogenous expression pattern of SNRK in Drosophila, we generated a knock-in Gal4 driver line, *Snrk-KI-Gal4*, using a PhiC31-mediated cassette exchange strategy designed to insert the gene encoding the yeast transcription factor Gal4 immediately downstream of the SNRK gene while preserving all endogenous regulatory elements (Figure S6A). Gal4 binding to the upstream activating sequence (UAS) led to green fluorescent protein (GFP) expressing and thus visualization of SNRK expression pattern (*Snrk-KI-Gal4>UAS-mCD8::GFP* flies in Figure S6B).

We generated a second targeted knock-in driver, *Snrk-KO-Gal4*, in which the SNRK coding sequence was replaced by Gal4 under the direct control of the SNRK promoter (Figure 5A). Confocal imaging of brains from flies with *Snrk-KO-Gal4 > UAS-mCD8::GFP* revealed an expression pattern (Figure 5B, Figure S6C). While there were differences between those two knock-in lies, both showed neuronal expression in regions such as the mushroom body (MB) including the calyx, MB medial lobes, and MB α lobe. Thus, SNRK is expressed in the nervous system.

**Figure 5.**
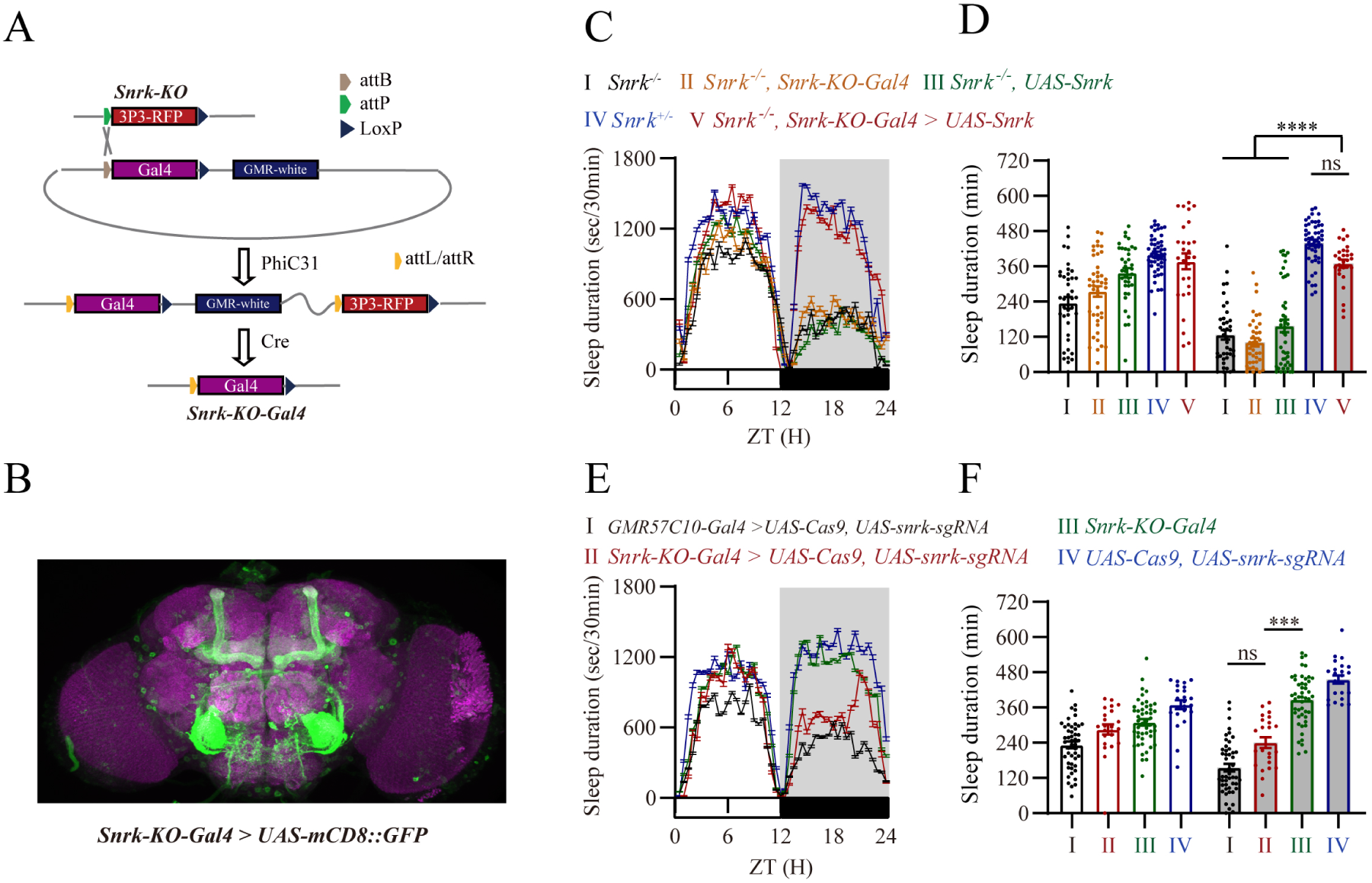
Necessity and Sufficiency of a Neuronal Subset Positive for *Snrk-KO-Gal4* in Sleep Regulation. **(A)** A schematic representation of the PhiC31 integrase-mediated cassette exchange strategy used to generate the targeted *Snrk-KO-Gal4* knock-in driver line. The 3P3-RFP cassette within the Snrk-KO locus was replaced with a Gal4 sequence under the direct control of the immediate SNRK promoter. **(B)** Representative confocal maximal projection images of an adult Drosophila brain expressing the membrane-targeted GFP reporter (Snrk-KO-Gal4 > UAS-mCD8::GFP). **(C, D)** Sleep profile **(C)** and quantification of daytime and night-time sleep duration (min) **(D)** for targeted spatial rescue experiments. **(E, F)** Sleep profile **(E)** and quantification of daytime and night-time sleep duration (min) ns, not significant; *p < 0.05; **p <0.01; ***p <0.001 and ****p <0.0001; mean ± standard error of the mean (mean ± SEM). Kruskal-Wallis test **(D, F).**

We functionally investigated whether the neuronal population labeled by the *Snrk-KO-Gal4* driver was sufficient in sleep regulation. Expression of wt SNRK exclusively in these neurons within the SNRK null background completely rescued the sleep loss and restored total sleep duration to wt control levels (Figure 5 C and D).

We validated the spatial requirement through CRISPR/Cas9 knockout strategy. Targeted disruption of SNRK restricted to the *Snrk-KO-Gal4* pattern (II: *Snrk-KO-Gal4>UAS-Cas9, UAS-snrk-sgRNA* in Figure 5 D and F) significantly reduced sleep duration. This localized knockout phenocopied the severe sleep loss observed in the pan-neuronal somatic knockout (I: *GMR57C10-Gal4>UAS-Cas9, UAS-snrk-sgRNA*). Both knockout groups exhibited reduced sleep compared to their respective transgenic controls (III: *UAS-Cas9, UAS-snrk-sgRNA*; IV: *GMR57C10-Gal4, UAS-Cas9*) (Figure 5 E and F).

Results from these bidirectional functional assays establish that the neurons captured by the *Snrk-KO-Gal4* driver are necessary and sufficient for SNRK regulation of sleep.

### Chemoconnectomic Analysis of SNRK Function in Sleep Regulation

Because SNRK functions in the nervous system, we decided to investigate the neurochemical features of neurons in which SNRK regulates sleep. CCTomics created by us is an efficient system to investigate neuronal requirement of any gene in *Drosophila*^55,56^.

For the present purpose of studying SNRK function, we used 223 lines of flies from CCTomics which expressed Gal4 in neurons expressing different neurotransmitters, their receptors, synthesizing enzymes or transporters^56^. Each CCT Gal4 was used to drive RNAi for SNRK before analysis of sleep. Normalizing the total sleep duration of each RNAi knockdown line to its respective genetic controls revealed a broad, continuous distribution of sleep phenotypes (Figure 6A; Table S1), ranging from severe sleep reduction to significant sleep enhancement.

**Figure 6.**
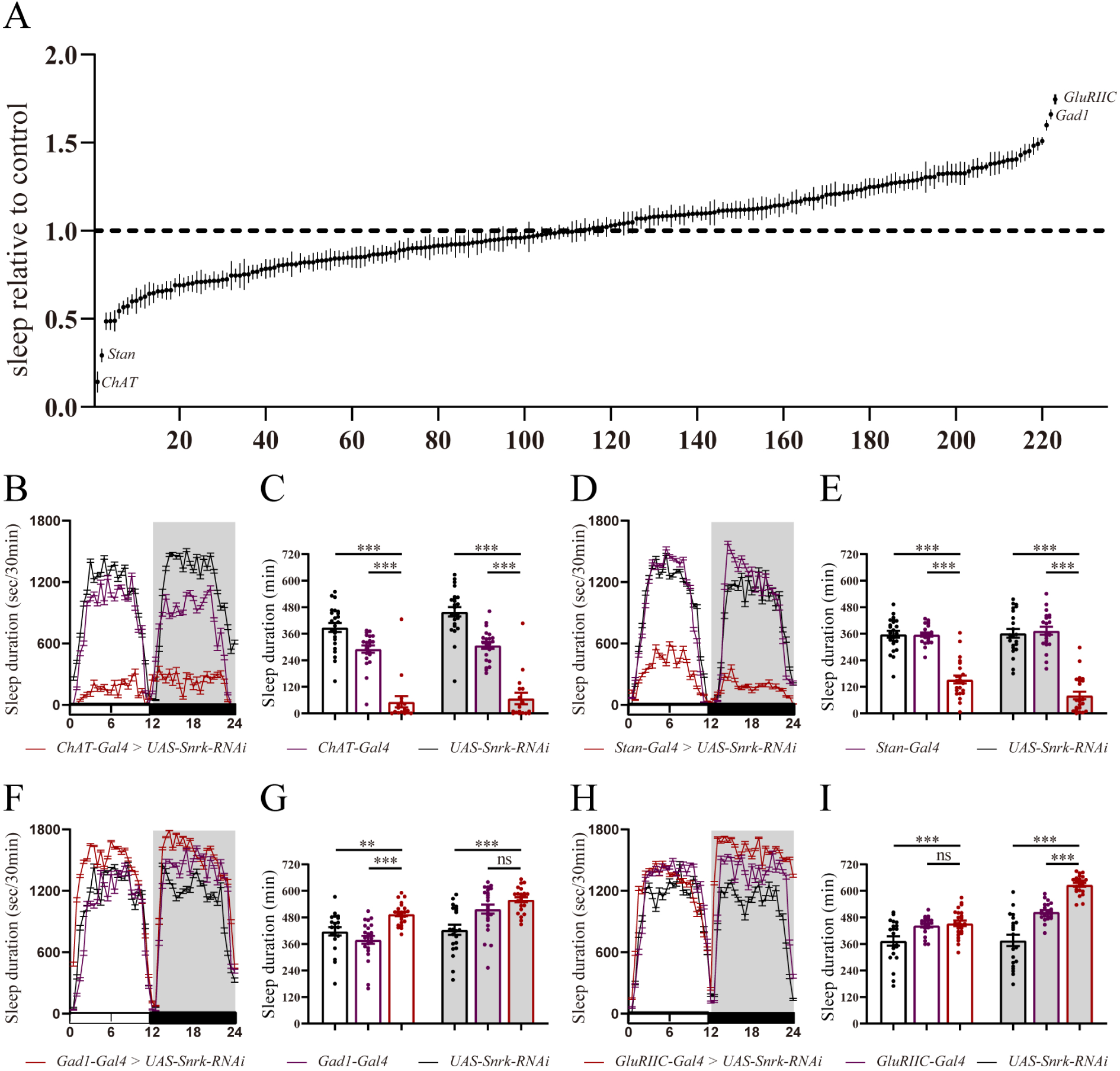
A CCT Screen of Neurons Required for SNRK Regulation of Sleep. **(A)** Distribution of normalized total sleep duration across a screening panel of 223 Gal4 driver lines crossed with *UAS-Snrk-RNAi*. The y-axis represents the total sleep duration of each knockdown line relative to its respective genetic controls. The dashed line (1.0) indicates the baseline control level. **(B, D)** Representative 24-hour sleep profiles (Sleep duration sec/30min) following targeted RNAi-mediated knockdown of SNRK in excitatory cholinergic neurons (*ChAT-Gal4*, **B**) and *Stan+* neuronal populations (*Stan-Gal4*, **D**), plotted alongside their corresponding *Gal4* and *UAS* transgenic controls. **(C, E)** Quantification of daytime and night-time sleep duration (min) for the *ChAT-Gal4* **(C)** and *Stan-Gal4* **(E)** knockdown experimental groups and their genetic controls. **(F, H)** Representative 24-hour sleep profiles (Sleep duration sec/30min) following targeted SNRK knockdown in inhibitory GABAergic neurons (*Gad1-Gal4*, **F**) and glutamatergic neurons (*GluRIIC-Gal4*, **H**). **(G, I)** Quantification of daytime and night-time sleep duration (min) for the *Gad1-Gal4* **(G)** and *GluRIIC-Gal4* **(I)** knockdown experimental groups and their genetic controls. ns, not significant; *p < 0.05; **p <0.01; ***p <0.001 and ****p <0.0001; mean ± standard error of the mean (mean ± SEM). Kruskal-Wallis test **(C, E, G, I).**

Targeted knockdown of SNRK in cholinergic neurons, with SNRK RNAi driven by ChAT-Gal4, drastically reduced total sleep duration and severely disrupted the daily sleep profile (Figure 6 B and C). Knockdown of RNAi in neurons labeled by Stan-Gal4 was also validated to reduce sleep significantly (Figure 6 D and E). To rule out potential off-target effects of RNA interference, we validated these requirements using an independent in vivo CRISPR/Cas9 somatic knockout approach. Targeted gene ablation of SNRK in ChAT+ populations reproduced the severe reduction in night-time and total sleep (Figure S7 A-D). Notably, the magnitude of sleep loss in these specific knockdown and knockout flies closely mimicked the severe insomnia phenotype of the SNRK null mutants.

Our screen also identified neuronal subsets in which SNRK RNAi yielded a contrasting physiological effect. Specific knockdown of SNRK in GABAergic neurons (Gad1-Gal4) or GluRIIC-Gal4 driven neurons resulted in a significant increase in total sleep duration compared to the controls (Figure 6 F-I). However, we did not follow these findings because SNRK null did not show such phenotypes. We therefore focus on ChAT+ neurons in which SNRK knockdown showed the strongest sleep loss phenotype among all CCT neurons.

### SNRK Antagonism of Cholinergic Inhibition of Sleep

To investigate structural and functional relationships between SNRK-expressing cells and the cholinergic system, we used an intersectional genetic reporter strategy with *Snrk-KO-Gal4* driven *UAS-Flp* and *ChAT-LexA* driven *LexAop-FRT-stop-FRT-GFP*. This allowed us to visualize neural populations expressing both SNRK and the cholinergic marker ChAT. Confocal imaging revealed distinct double labeling, indicating endogenous SNRK expression in defined subsets of cholinergic neurons (Figure 7I).

**Figure 7.**
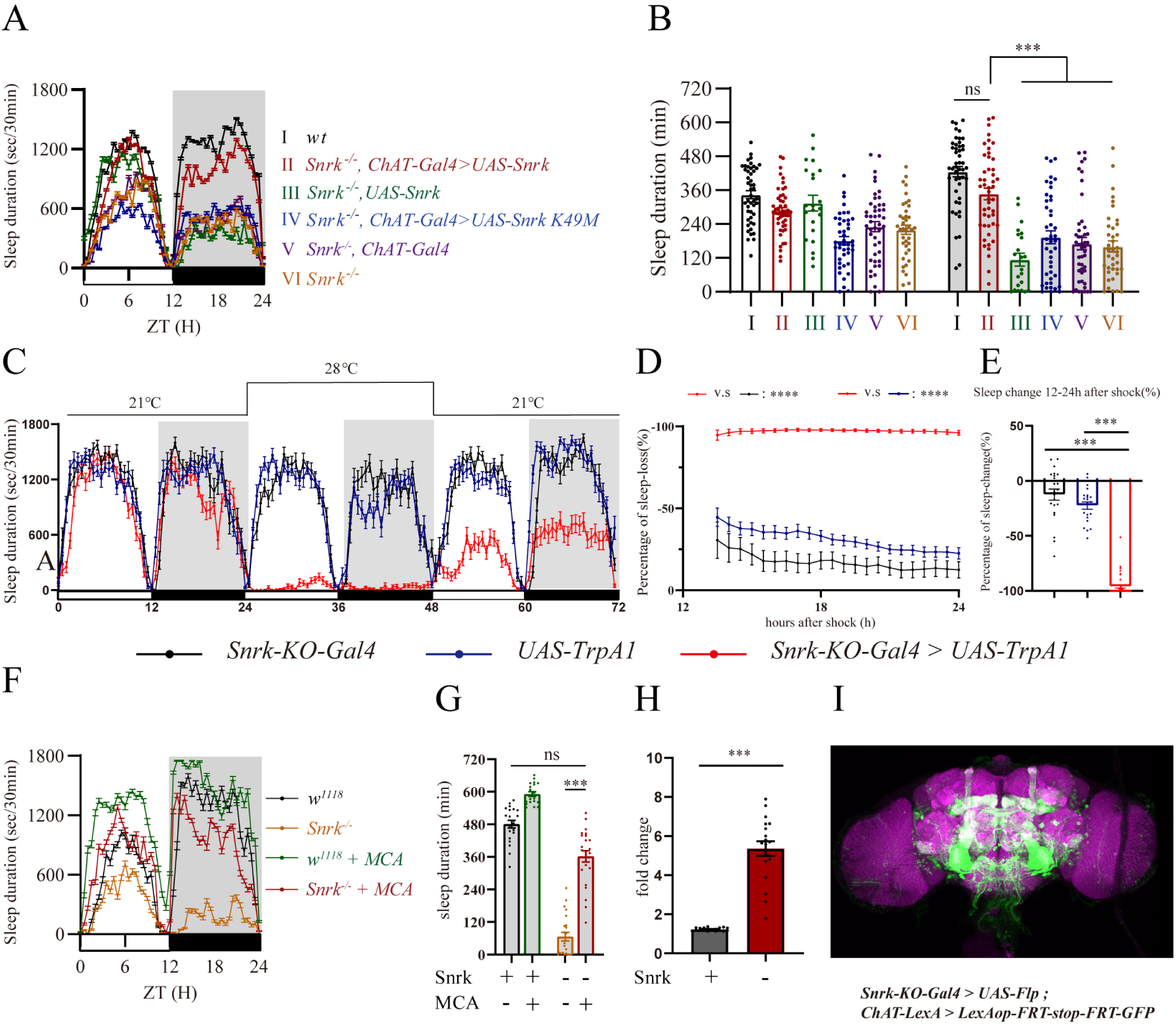
Functioning of SNRK in Cholinergic Neurons. **(A)** Expression of either wild-type SNRK (*UAS-Snrk*) in the SNRK null mutant background, plotted alongside respective genetic controls. **(B)** Quantification of daytime and night-time sleep duration (min) for in (A), along with the expression of the kinase-dead variant (*UAS-Snrk-KM*) specifically within cholinergic neurons (*ChAT-Gal4*). **(C)** Continuous 72-hour sleep profile during thermogenetic activation of *Snrk*-expressing neurons. Experimental flies (*Snrk-KO-Gal4 > UAS-TrpA1*) and controls were monitored at a baseline permissive temperature (21 °C), shifted to a restrictive temperature (28 °C) to induce TrpA1-mediated depolarization, and subsequently returned to 21 °C. **(D)** Hourly tracking of the percentage of sleep loss (%) during the 12-hour restrictive temperature window (hours 14 to 24 after the temperature shock). **(E)** Quantification of the overall sleep change (%) during the 12-24 hour period following the shift to 28 °C, normalized to baseline sleep levels. **(F)** Continuous 48-hour sleep profile of wild-type (*w1118*) controls and SNRK null mutants administered either vehicle or the specific nicotinic acetylcholine receptor (nAChR) antagonist mecamylamine (MCA). **(G)** Quantification of nighttime sleep duration (min) for the pharmacological MCA treatment groups and their respective untreated controls. **(H)** Fold change in sleep duration induced by MCA treatment relative to the untreated baseline for wild-type controls (+) and SNRK null mutants (-). **(I)** Representative confocal maximal projection image of an adult *Drosophila* central brain utilizing an intersectional genetic reporter strategy. Green fluorescence labels neuronal somata and projections that co-express both SNRK (*Snrk-KO-Gal4 > UAS-Flp*) and the cholinergic marker *ChAT* (*ChAT-LexA > LexAop-FRT-stop-FRT-GFP*). ns, not significant; *p < 0.05; **p <0.01; ***p <0.001 and ****p <0.0001; mean ± standard error of the mean (mean ± SEM). Kruskal-Wallis test **(B, E, G)**; Unpaired t-test **(H).**

We then tested whether the enzymatic function of SNRK within these cholinergic neurons was sufficient to restore sleep. Targeted expression of wt SNRK utilizing the *ChAT-Gal4* driver in the SNRK mutant background significantly rescued the severe reduction in total sleep duration and restored the 24-hour sleep profile (Figure 7 A and B). Cholinergic-specific wt SNRK complementation completely normalized both sleep bout duration (Figure S8A) and sleep bout number (Figure S8B), rendering them indistinguishable from controls. Expression of the kinase-dead variant (*UAS-Snrk-KM*) driven by the same cholinergic promoter failed to rescue the total sleep defects or the severe sleep fragmentation (Figure 7 A and B, Figure S8 A and B).

Cholinergic signaling in the Drosophila central nervous system is well known to cause wakefulness and inhibit sleep^57,59,66^. The finding of SNRK function in cholinergic neurons immediately led to the hypothesis that the physiological role of SNRK is to inhibit cholinergic neurons.

To test this, we activated the SNRK-expressing neurons with TrpA1 (*Snrk-KO-Gal4 > UAS-TrpA1*)^67,68^. Upon shifting the environmental temperature to 28°C to activate the TrpA1 channels, the experimental flies exhibited a near-total loss of sleep (Figure 7 C, D and E), confirming that activation of the SNRK expressing neurons was similar to activation of cholinergic neurons in their inhibition of sleep.

If the extreme sleep loss observed in SNRK null mutants is a direct consequence of hyperactive cholinergic transmission, then pharmacological blockade of downstream acetylcholine receptors should reverse the mutant phenotype. To further test this model, we treated adult flies with mecamylamine (MCA), a specific nicotinic acetylcholine receptor (nAChR) antagonist^58,69^ (Figure 7F). While continuous MCA administration had moderate effect on baseline sleep duration of wt flies, it robustly rescued the night-time sleep loss with the bout number and duration in SNRK mutants (Figure 7 G and H, Figure S9 A and B). Quantification revealed a highly significant increase in sleep duration in the MCA-treated mutant group.

## DISCUSSION

We have obtained results from genetic, intersectional, thermogenetic and pharmacological experiments which establish an important role for SNRK in regulating sleep, and suggest a mechanistic model: SNRK is expressed in a subset of wake-promoting cholinergic neurons to inhibit cholinergic signaling, thereby promoting sleep.

### Essential Role of SNRK in Sleep

The LOF sleep phenotype of SNRK observed here in flies is stronger than those of all the other kinases in flies or mice^14,16,18,24,26,27,30,34,36,70–73^. While sleep was dramatically increased in GOF SIK mutants^38–40^, there was no sleep phenotype in SIK1 and SIK2 LOF mutants^45^, and only moderate sleep changes in SIK3 LOF mutants^14,16,45^. The much stronger LOF sleep phenotype of SNRK suggests this member of the ARK family as an important regulator of sleep.

SIK 1, 2 and 3 are also ARKs, it is unclear whether their GOF phenotypes were physiological, or mimicking the functions of other kinases. If it was the latter case, SNRK would be a natural candidate as an ARK which could potentially be mimicked by the SIKs. In any case, the physiological role of each SIK in sleep is not as strong as SNRK, at least in flies.

The kinase activity is strictly required for SNRK to regulate sleep. The re-introduction of the kinase-dead mutant K49M completely failed to rescue the insomnia of null mutants, and the phospho-resistant T177A mutant exhibited a significantly impaired rescue capacity. This absolute catalytic dependence aligns SNRK with other known sleep-regulatory kinases^14,32,38,74^, which function through dynamic phosphorylation rather than structural roles.

It is important to distinguish whether a gene affect sleep because of its role in development (e.g., *inc*, *Cul3*, or *Fmr1*)^75–78^ or adulthood. SNRK acts in adulthood to regulate sleep because depletion of SNRK in adulthood was sufficient to induce severe sleep loss, whereas adult-specific restoration effectively reversed insomnia of null mutants. Together, these findings demonstrate SNRK can regulate sleep independently of early development, acting as an active requirement in adulthood.

### Cholinergic Requirement for SNRK Regulation of Sleep

A powerful approach is CCT^55,56^. Our unbiased CCT screen mapped the functional requirement of SNRK across 223 diverse neurochemical subpopulations.

Expression of SNRK RNAi in ChAT+ neurons yielded the strongest sleep loss phenotype. Conditional knockout of SNRK in ChAT+ neurons using CRISPR/Cas9 double-confirmed the strong sleep loss. Restoration of SNRK within ChAT+ neurons in the SNRK null background was sufficient to rescue sleep loss, demonstrating SNRK is required within the cholinergic neurons to regulate sleep.

Cholinergic transmission in *Drosophila* is wake-promoting^57,59,66^. Both the SNRK null mutants and the TrpA1 activation in *SNRK-KO-Gal4* driven neurons phenocopied insomnia caused by hyperactivation of ChAT+ neurons. These results support that SNRK inhibits the cholinergic signaling, and that loss of SNRK leads to cholinergic hyperactivity. Our initial attempts in manipulation of ChAT+ neurons using TrpA1 activation or Shibire^ts^ silencing within the SNRK null background resulted in lethality (Figure S10 A and B). Therefore, the pharmacological intervention was used to support the mechanistic hypothesis: blocking nicotinic acetylcholine receptors (nAChRs) with MCA successfully restored sleep in SNRK null mutants by suppressing the cholinergic signaling.

The precise downstream molecular substrates of SNRK remain unidentified. Future studies aimed at systematically identifying SNRK-dependent phosphorylation events will be important for elucidating its molecular function, bridging the gap between this kinase’s biochemical activity and its behavioral output.

## METHODS

### Fly stocks

Flies were reared on standard corn meal at 25 ℃, 60% humidity with a 12 h : 12 h light : dark photoperiod. UAS-Cas9 and sgRNAs lines were generated in our laboratory. Flies for regional-screen were from the Bloomington Stock and CCT-screen were from our laboratory. All flies used in behavioral assays were backcrossed for at least 7 generations. All results of sleep analysis in this paper were obtained from male flies.

### Generation of KO, KI and transgenic fly lines

The cDNA of CG8485 was cloned by reverse transcription of mRNA extracted from w1118 flies. The coding sequence of CG8485 was then constructed into pACU2 plasmid as pACU2-UAS-CG8485-WT. Point mutation at K49M, T177A, T177E, S586D and S586A was done by Gibson assembly based on pACU2-UAS-CG8485-WT. The plasmids were then injected into attp40 site to obtain relevant fly strains.

KO and KI lines were generated as described previously^56^. KO flies were generated with the CRISPR/Cas9 system. Two different sgRNAs were constructed with U6b-sgRNA plasmids to introduce two double strand break (DSB). 5’ homologous arm and the 3’ homologous arm of ∼2 kb amplified from fly genome were inserted into a pBSK plasmid for homologous recombination repair. The cassette of attP-3P3-RFP was introduced in the middle. sgRNA plasmids and the donor plasmids were injected into vas-Cas9 embryos to introduce attP-3P3-RFP into the genome at the region of interest and replaced it by homologous recombination. 3P3-RFP served as a marker to screen for the correct flies. Primers across the homologous arms were designed to verify the sequences by PCR and DNA sequencing. attP site was introduced into the genome with 3P3-RFP-LoxP. For KI flies, the nos-phiC31 virgin females were first crossed with knock-out males and the pBSK plasmid inserted with attB-T2A-Gal4-miniwhite-LoxP cassette was injected into the female embryos. Miniwhite serves as a marker to screen for the correct flies, which could be excised by the Cre/LoxP system. Primers were designed to verify the sequence by PCR and DNA sequencing.

### Quantitative Real-time PCR

The mRNA of flies were extracted at the age of 5∼7 days or after behavior assays by TRIzol reagent. The genomic DNA was removed and the mRNA were reverse transcribed by cDNA Synthesis Kit. Quantitative PCR was carried out by with TransStart Top Green qPCR SuperMix kit (TransGen) in the Bio-Rad PCR system (CFX96 Touch Deep Well). The sequences of primers for detection are:

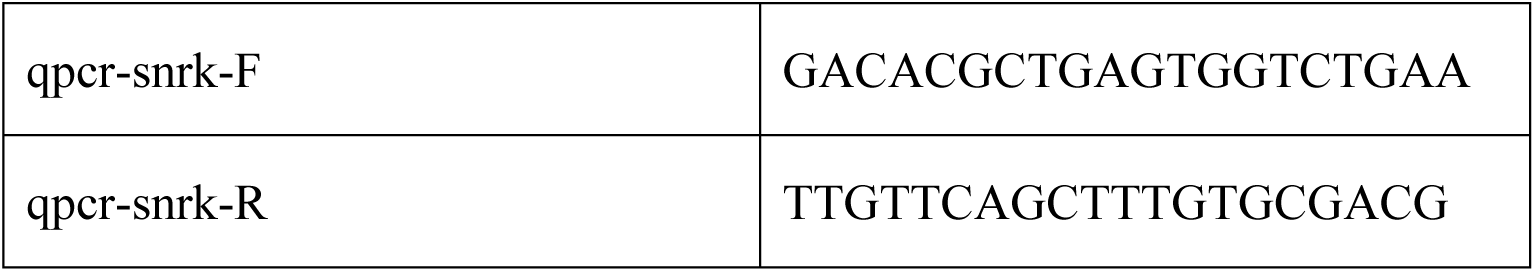
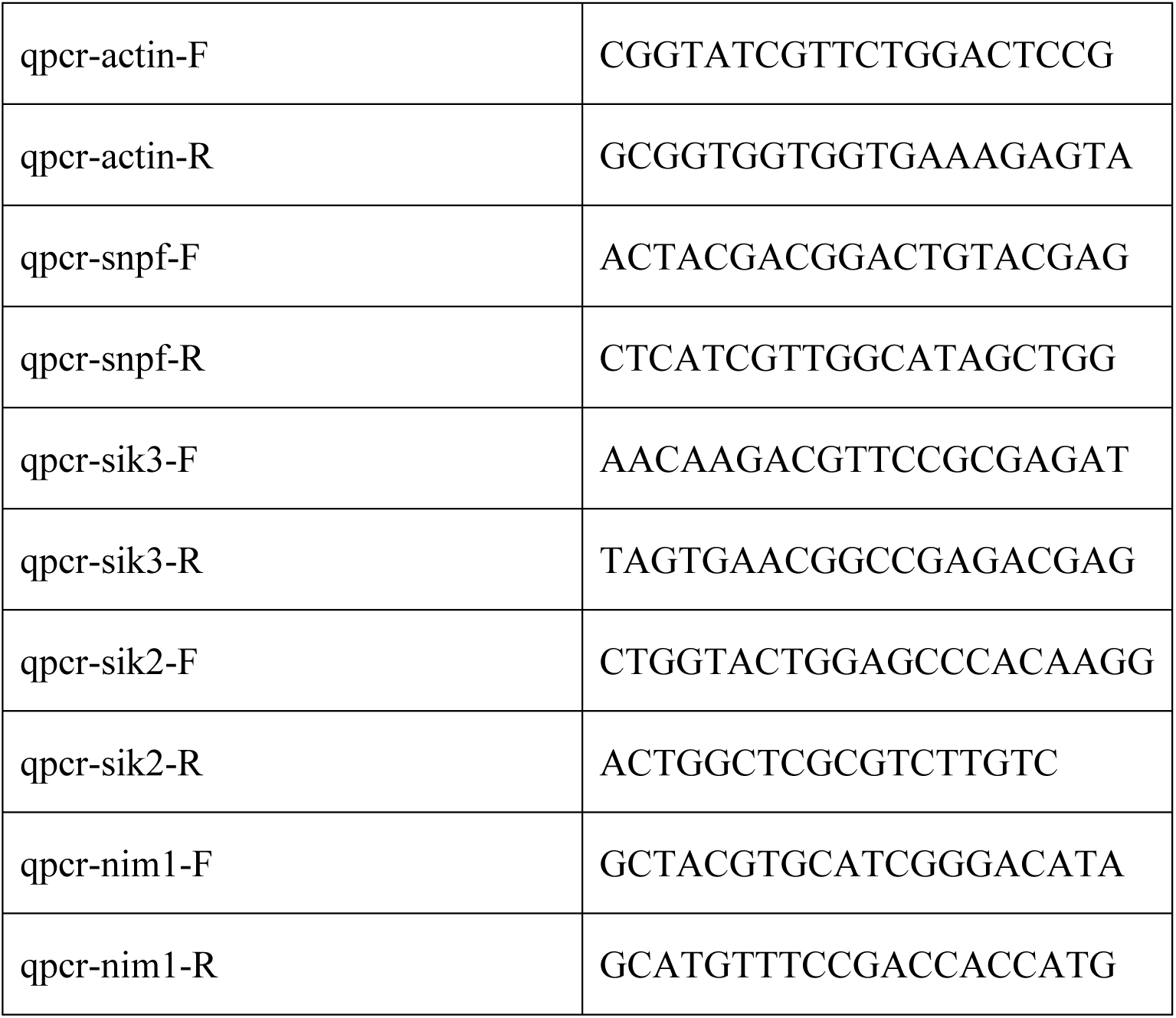

### Drosophila sleep analysis

Drosophila sleep analysis was performed as described previously^79,80^. 5–7 days old flies were placed in a 65 mm # 5 mm clear glass tube with one end containing food and the other end plugging with cotton. All flies were recorded by video-cameras. Before data acquisition, flies were entrained to an LD cycle at 25 ℃, 50% humidity for at least 2 days with infrared LED light providing constant illumination when lights off. Immobility longer than 5 min was defined as one sleep event^49,81^. Information of fly location was tracked and sleep parameters were analyzed using Matlab (Mathworks), from which dead flies were removed.12-48 flies were recorded for each group and data was shown as individual flies for every bar graph. Sleep duration, sleep bout duration, sleep bout number, and sleep latency for each LD were analyzed.

### Drosophila circadian analysis

Flies were reared and recorded in the same condition as sleep assay as described in papers from our laboratory^57,80^, except that the condition was constant darkness. Six to eight days activity was measured and calculated in ActogramJ^82^. Rhythmic strength, power and period were calculated by Chi-square method.

### Conditional gene manipulation at adult stage

Flies for gene manipulation at adult stage were raised on standard corn meal until 5∼7 days old. For gene expression, a diet of 1 mM Ru486 added into the corn meal was prepared for induction. Same amount of ethanol was added into the corn meal for the control group. Flies were switched to corresponding diet and fed *ad libtum* for 2 days before video recording. For gene knockdown, parental flies were crossed and the selected offsprings were raised at 22 ℃ before video recording. Videos of 3 days at 22 ℃ was recorded as the baseline and 1 day at 29 ℃ as activation, followed by 3 days at 22 ℃ as recovery. Percentage of sleep-loss was calculated by comparing the sleep duration at certain time point with that of 24 h before. Other procedures was identical to the sleep analysis described above.

### MCA administration

For the treatment of nicotinic acetylcholine receptor inhibitor mecamylamine hydrochloride (HY-B1395, MedChemExpress), a 100 mM stock solution was diluted to the desired concentration in a food mixture containing 5% sucrose and 2% agar, which had been boiled and then cooled to below 70 °C. Other procedures was identical to the sleep analysis described above.

### The CCT screen

The transgenic lines were adapted from the previous paper of our laboratory. For CCT lines from different genetic background (*w1118* or *IsoCS*), the control group was set by crossing UAS-Snrk-RNAi with either *w1118* or *IsoCS* in every batch. Sleep duration relative to control was analyzed by comparing the sleep duration of the screened CCT lines with the corresponding control group.

### Neuronal excitability manipulation

Parental generation flies were crossed and placed under 18℃ conditions. Males were selected and maintained at 18 ℃ for 8 to 10 days and then recorded at 21 ℃ for 3 days as the baseline and at 29 ℃ for 1 day activation followed by 1 day recovery. Percentage of sleep-loss was calculated by comparing the sleep duration at certain time point with that of 24 h before.

### Immunostaining and confocal imaging

For all immunostainings, 6 to 10 days old adult flies were anesthetized and dissected in ice-cold phosphate buffered saline (PBS). Whole-mount brains were fixed in 2% paraformaldehyde (wt/vol) for 55 min at room temperature, washed three times in washing buffer (PBS containing 3% sodium chloride and 1% Triton X-100 (vol/vol)) for 15 min. Brains were subject to 10% normal goat serum (diluted in 2% PBST) overnight at 4 ℃, before incubation with the primary antibody in dilution buffer (1% normal goat serum in 0.25% PBST) for 24 h at 4 ℃. Samples were washed for three times for 15 min before incubation with the secondary antibody in the dilution buffer for 24 h in darkness at 4 ℃. Samples were washed three times for 15 min, before being mounted on slices in Focus Clear (Cell Explorer Labs, FC-101), and visualized on a Zeiss LSM 710 confocal microscope. Images were processed by Imaris, ZEN blue and Arivis pro softwares.

### Statistics

All statistical analyses were performed with Prism 7 (GraphPad Software). Samples larger than 2 groups were analyzed using Kruskal-Wallis test. Unpaired t-test or Mann-Whitley was used for 2 groups comparison. Power spectrum between different lines was compared by 2-way ANOVA followed by Turkey’s multiple comparisons test. n.s. denotes P > 0.05; *P< 0.05; **P < 0.01; and ***P < 0.001 for all statistical results in this paper.

---

**Figure S1.**
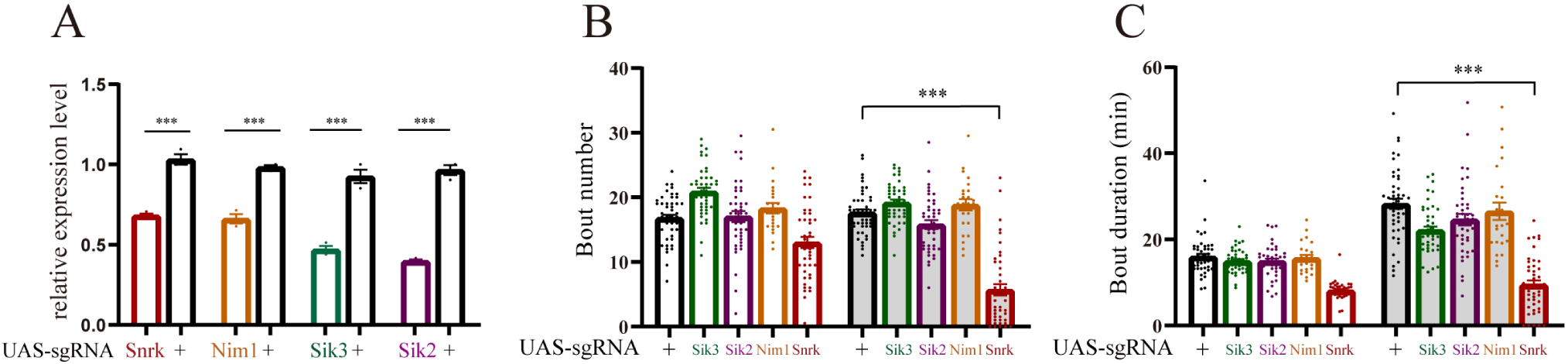
Detailed Sleep Analysis of Pan-neuronal CRISPR/Cas9 Screen of the ARKs. **(A)** Relative mRNA expression levels of the targeted genes (*Snrk*, *Nim1*, *Sik3*, *Sik2*) in the somatic knockout brains compared to corresponding controls. **(B, C)** Quantification of daytime and night-time sleep bout number (min) **(B)** and sleep bout duration **(C)**. ns, not significant; *p < 0.05; **p <0.01; ***p <0.001 and ****p <0.0001; mean ± standard error of the mean (mean ± SEM). Kruskal-Wallis test **(B, C)**; Unpaired t-test **(A).**

**Figure S2.**
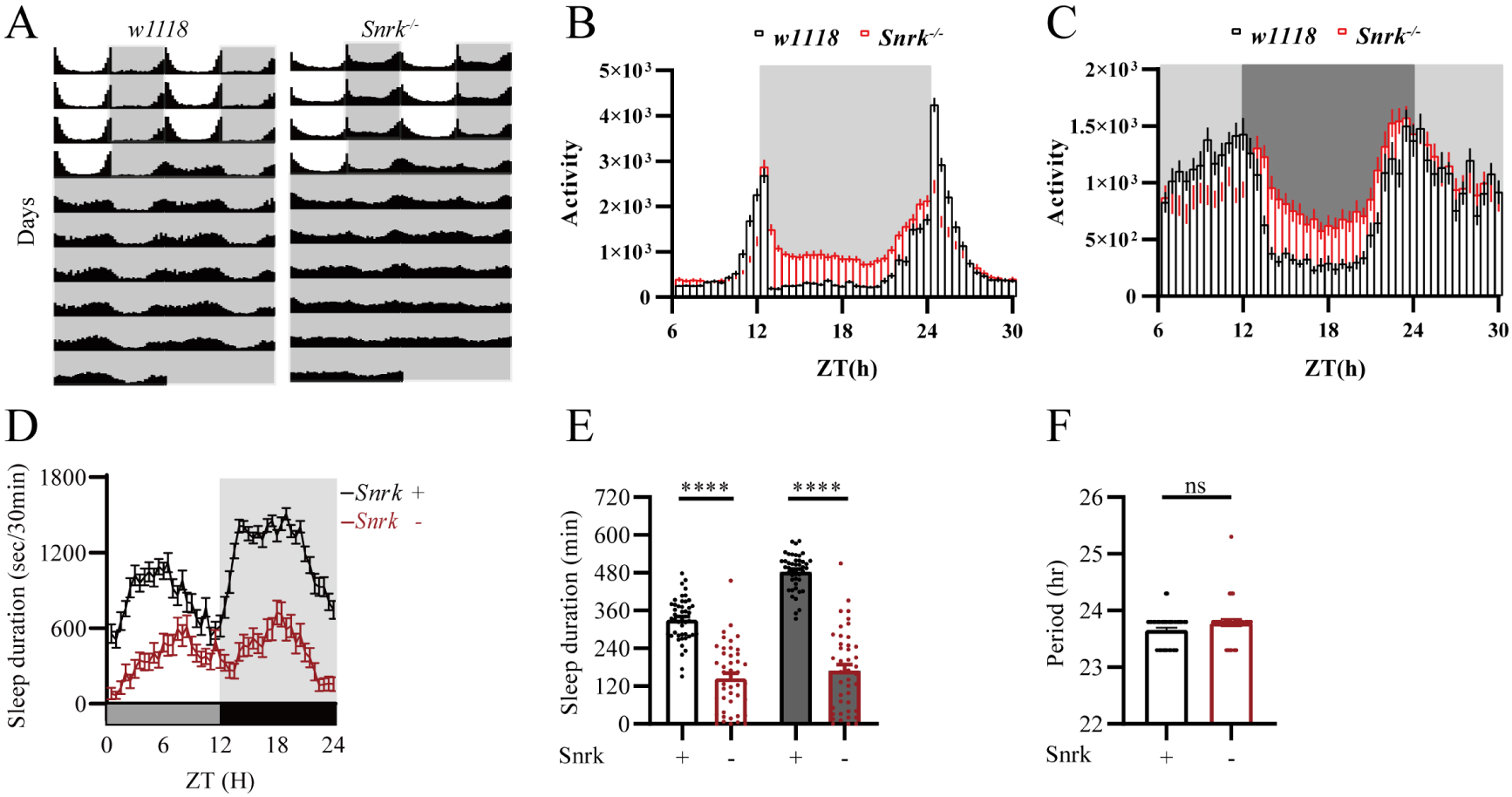
Assessment of Circadian Rhythm and Sleep in SNRK Mutants under Constant Darkness. **(A)** Representative double-plotted actograms of locomotor activity for wild-type (*w1118*) controls and SNRK null mutants (*Snrk^-/-^*) recorded under free-running constant darkness (DD) conditions for consecutive days. **(B, C)** Average daily locomotor activity profiles of the indicated genotypes during the DD period. **(D)** Continuous 24-hour sleep profile of control (*Snrk +*) and SNRK null mutants (*Snrk-*) monitored under constant darkness. **(E)** Quantification of total subjective daytime and subjective night-time sleep duration (min) under DD conditions. **(F)** Quantification of free-running circadian period under DD (hr) ns, not significant; *p < 0.05; **p <0.01; ***p <0.001 and ****p <0.0001; mean ± standard error of the mean (mean ± SEM). Mann-Whitney test **(E, F).**

**Figure S3.**
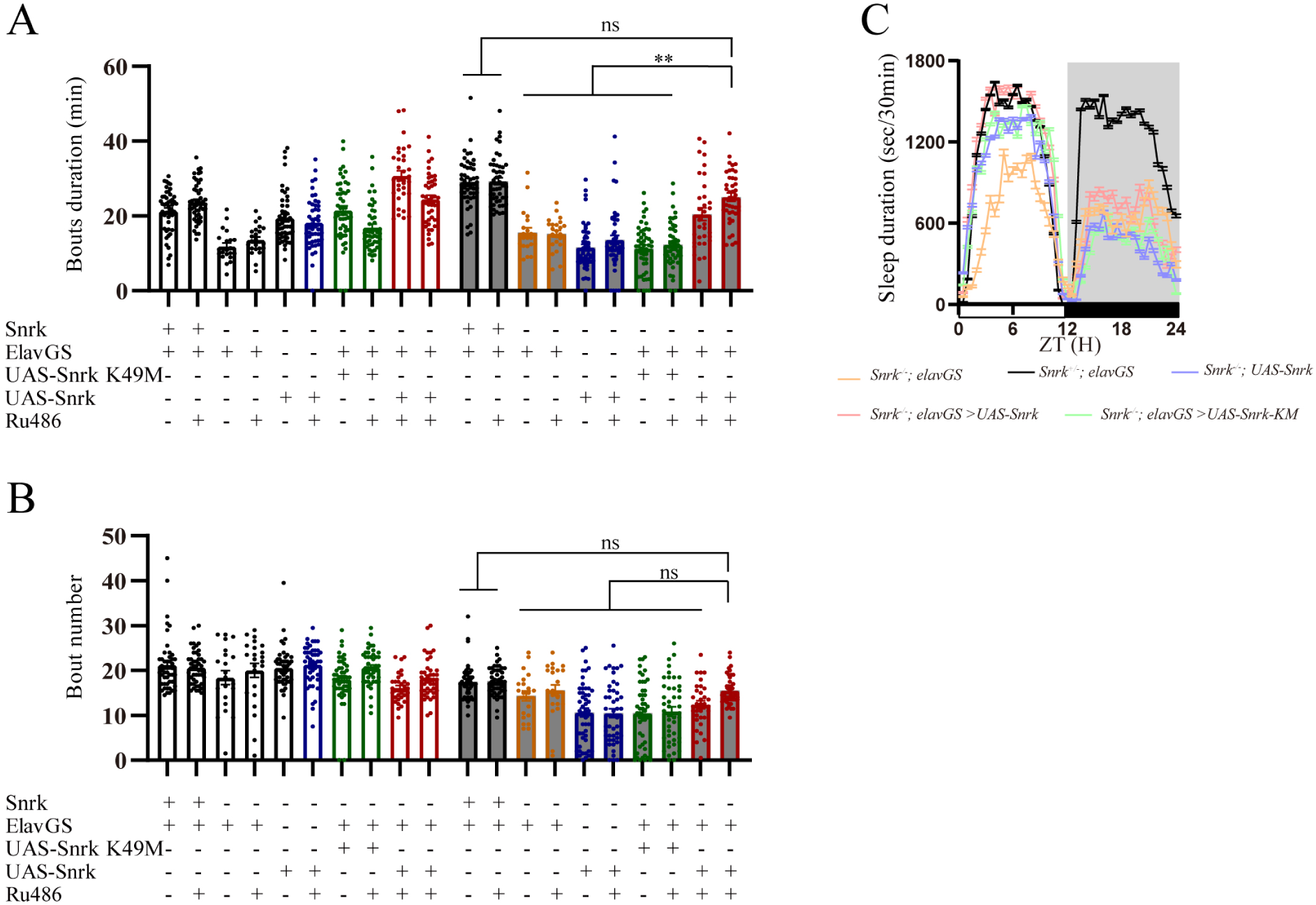
Detailed Sleep Analysis of Adult-Specific, Temporally Controlled SNRK Expression. **(A)** Quantification of sleep bout duration (min) for SNRK null mutants expressing either wild-type SNRK (*UAS-Snrk*) or the kinase-dead variant (*UAS-Snrk K49M*) pan-neuronally via the RU486-inducible GeneSwitch system (*elavGS*), alongside respective controls. **(B)** Representative 24-hour sleep profile of the experimental and control groups in the uninduced baseline state (without RU486 feeding). **(C)** Quantification of sleep bout number for the indicated genotypes in the presence (+) or absence (-) of the RU486 inducer. ns, not significant; *p < 0.05; **p <0.01; ***p <0.001 and ****p <0.0001; mean ± standard error of the mean (mean ± SEM). Kruskal-Wallis test **(A, B).**

**Figure S4.**
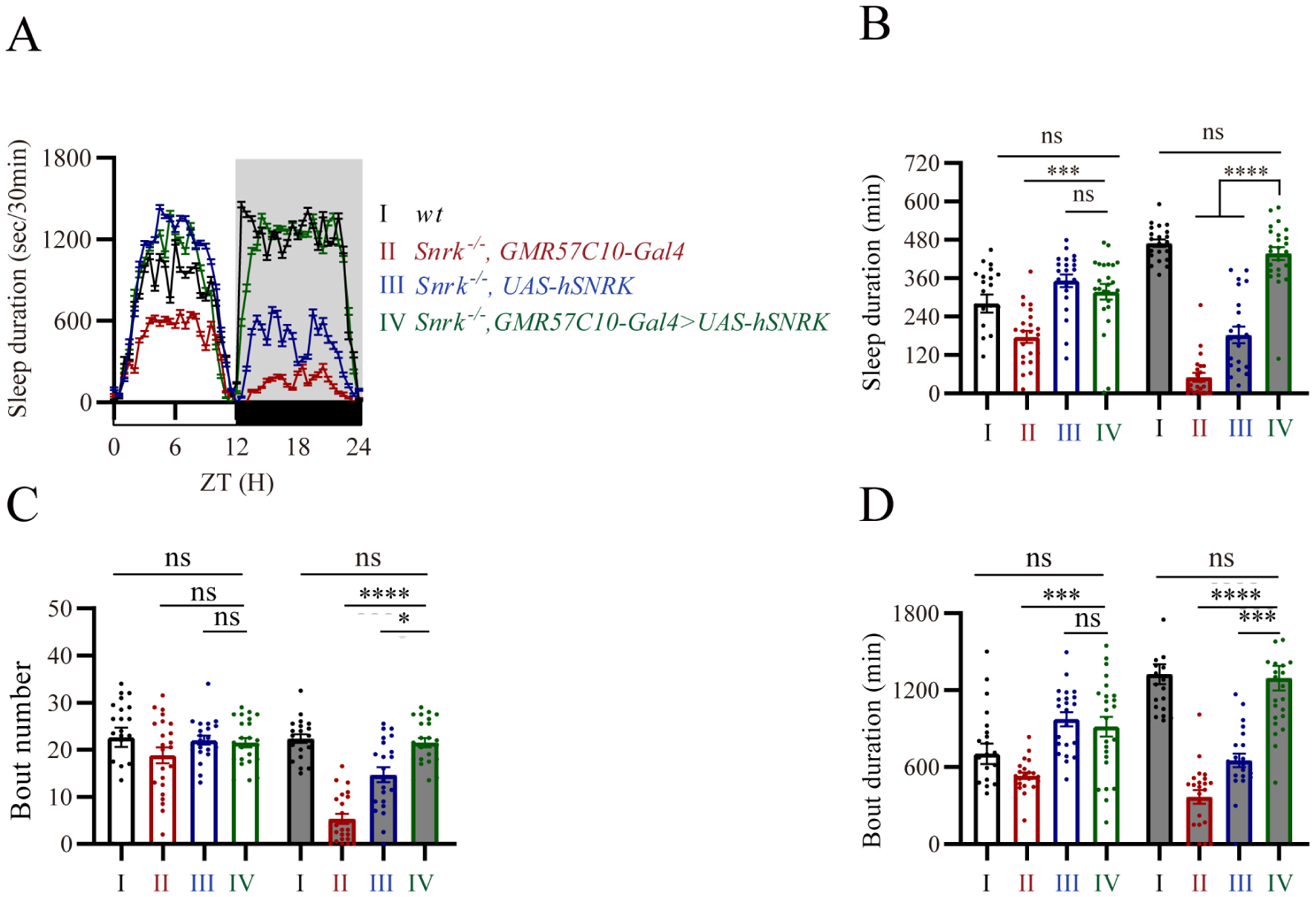
Pan-neuronal Expression of the Human SNRK Ortholog (hSNRK) in *Drosophila* SNRK mutants. **(A)** Quantification of daytime and night-time sleep duration (min) in SNRK null mutants expressing the human *SNRK* gene (*UAS-hSNRK*) pan-neuronally (*GMR57C10-Gal4*), plotted alongside corresponding genetic controls. **(B)** Continuous 48-hour sleep profile of the hSNRK transgenic rescue flies and their respective controls. **(C, D)** Detailed quantitative analysis of sleep architecture parameters, including sleep bout number **(C)** and sleep bout duration (min) **(D)**, for the indicated hSNRK rescue and control genotypes. ns, not significant; *p < 0.05; **p <0.01; ***p <0.001 and ****p <0.0001; mean ± standard error of the mean (mean ± SEM). Kruskal-Wallis test **(A, C, D)**

**Figure S5.**
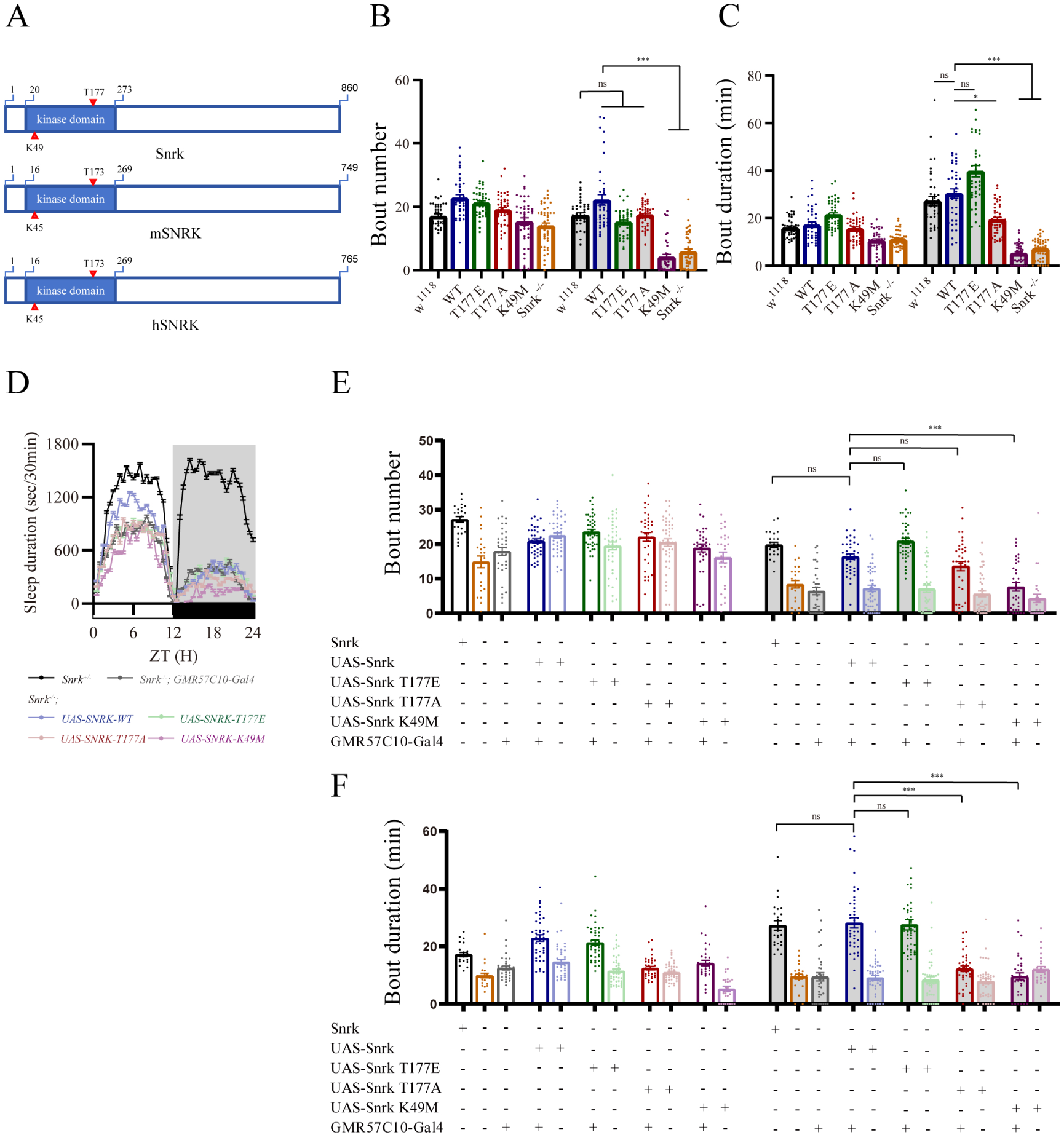
Detailed Sleep Analysis of SNRK knock-in and transgenic lines. **(A)** A schematic sequence alignment of the N-terminal kinase domains of *Drosophila* Snrk, mouse SNRK (mSNRK), and human SNRK (hSNRK). The phosphorylation target (threonine, T177 in *Drosophila*) and the critical ATP-binding residue (lysine, K49 in *Drosophila*) within the catalytic domain are indicated. **(B, C)** Quantification of daytime and night-time sleep bout number **(B)** and sleep bout duration (min) **(C)** for endogenous *Snrk* knock-in variants (WT, T177E, T177A, K49M), plotted alongside wild-type (*w1118*) and complete SNRK null (*Snrk ^-/-^*) controls. **(D)** 24-hour sleep profile of the control groups without GMR57C10-Gal4. **(E, F)** Quantification of daytime and night-time sleep bout number **(E)** and sleep bout duration (min) **(F)** for pan-neuronal transgenic rescue flies. ns, not significant; *p < 0.05; **p <0.01; ***p <0.001 and ****p <0.0001; mean ± standard error of the mean (mean ± SEM). Kruskal-Wallis test **(B, C, E, F).**

**Figure S6.**
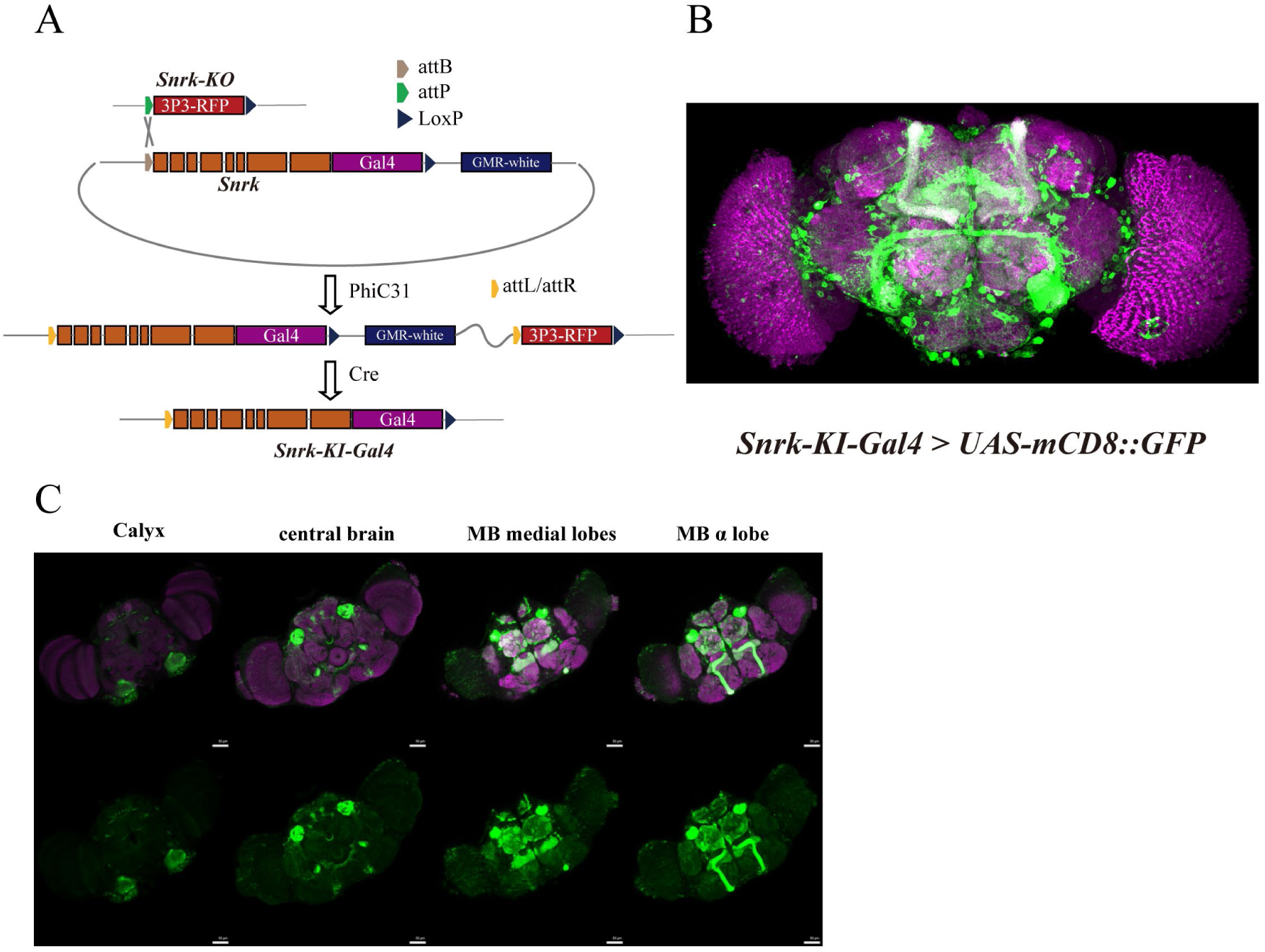
Expression Pattern of SNRK. **(A)** A schematic representation of the PhiC31 integrase-mediated cassette exchange strategy used to generate the *Snrk-KI-Gal4* driver line. The 3P3-RFP cassette within the Snrk-KO locus was replaced with the *Snrk* gene followed by a Gal4 sequence. **(B)** Representative confocal maximal projection image of an adult *Drosophila* brain showing the expression pattern of the *Snrk-KI-Gal4* driver (*Snrk-KI-Gal4 > UAS-mCD8::GFP*). **(C)** Detailed confocal maximal projection images of an adult *Drosophila* brain expressing the membrane-targeted GFP reporter (*Snrk-KO-Gal4 > UAS-mCD8::GFP*).

**Figure S7.**
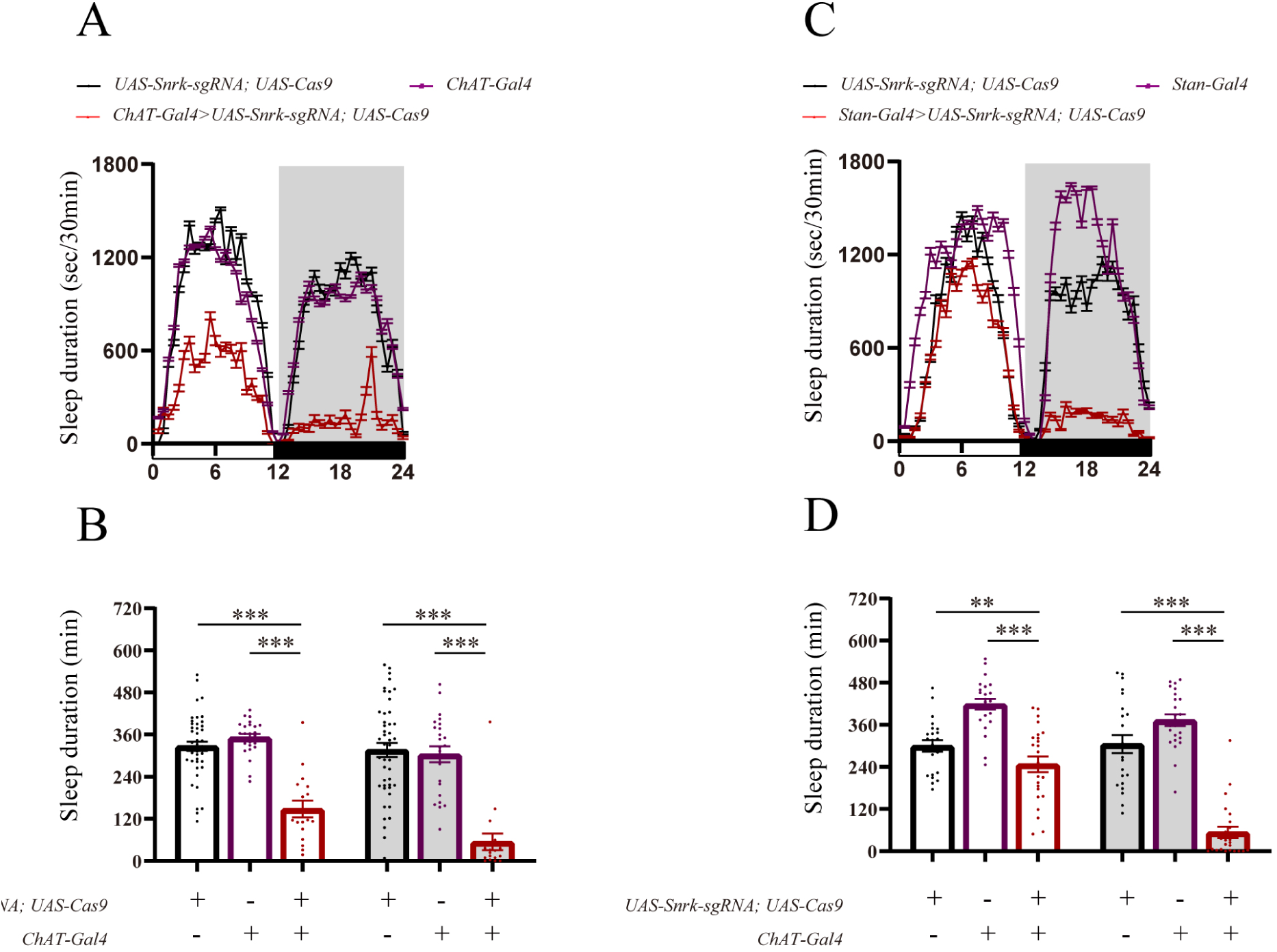
Targeted SNRK Knockout in ChAT and Stan Positive Neurons. **(A, B)** Continuous 24-hour sleep profile **(A)** and quantification of daytime and night-time sleep duration (min) **(B)** following knockout of SNRK within cholinergic neurons (*ChAT-Gal4 > UAS-Snrk-sgRNA, UAS-Cas9*), plotted alongside respective transgenic controls. **(C, D)** Continuous 24-hour sleep profile (Sleep duration sec/30min) **(C)** and quantification of daytime and night-time sleep duration (min) **(D)** following knockout of SNRK in Stan+ neuronal populations (*Stan-Gal4 > UAS-Snrk-sgRNA, UAS-Cas9*), plotted alongside respective transgenic controls. ns, not significant; *p < 0.05; **p <0.01; ***p <0.001 and ****p <0.0001; mean ± standard error of the mean (mean ± SEM). Kruskal-Wallis test **(B, D).**

**Figure S8.**
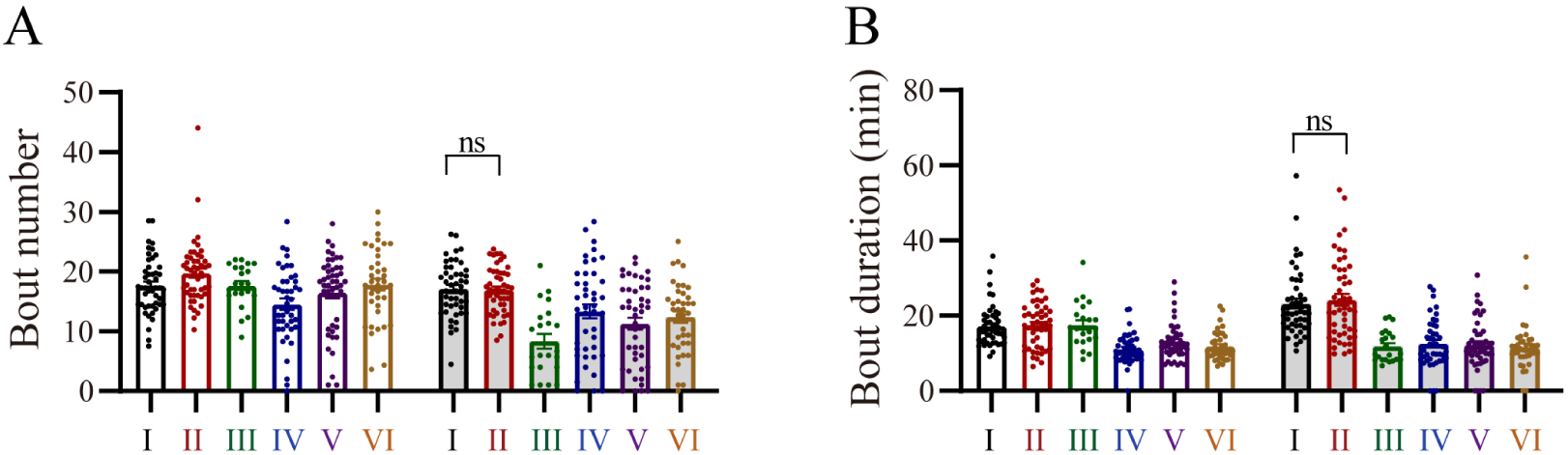
Detailed Sleep Analysis of SNRK Expressed in Cholinergic Neurons. **(A, B)** Quantification of daytime and night-time sleep bout number (min) **(A)** and sleep bout duration **(B)**. ns, not significant; *p < 0.05; **p <0.01; ***p <0.001 and ****p <0.0001; mean ± standard error of the mean (mean ± SEM). Kruskal-Wallis test **(A, B).**

**Figure S9.**
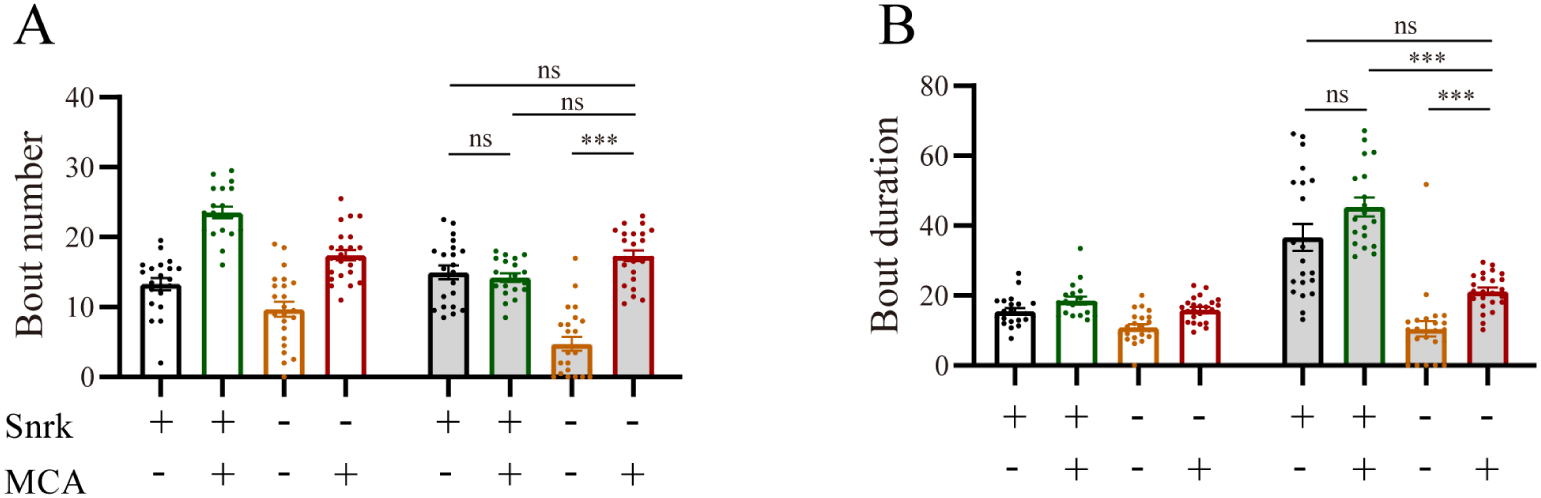
Detailed Sleep Analysis in the MCA Experiment. **(A, B)** Quantification of daytime and night-time sleep bout number (min) **(A)** and sleep bout duration **(B)**. ns, not significant; *p < 0.05; **p <0.01; ***p <0.001 and ****p <0.0001; mean ± standard error of the mean (mean ± SEM). Kruskal-Wallis test **(A, B).**

**Figure S10.**
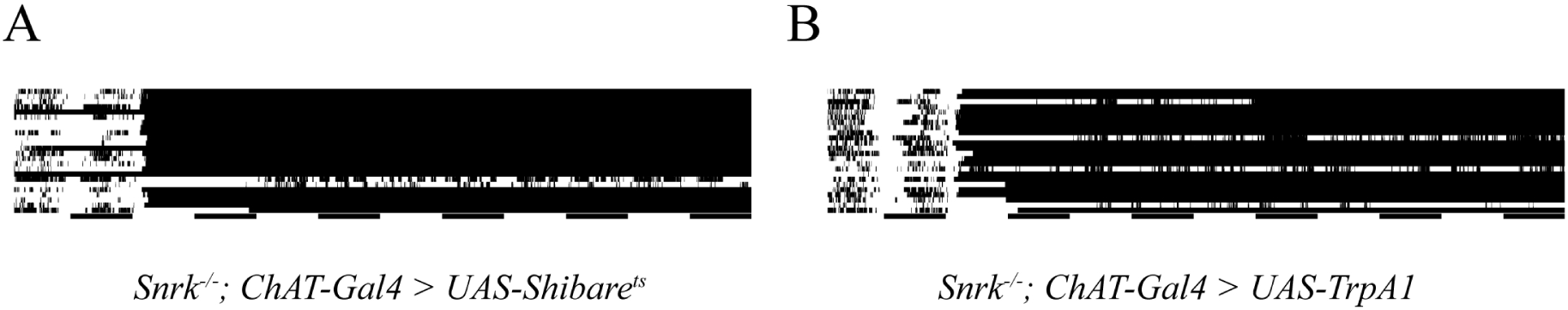
Actogram of ChAT Inhibition and Activation in SNRK null Mutants. **(A, B)** Actogram of ChAT+ neuron inhibition with UAS-Shibare^ts^ (**A**) and activation with UAS-TrpA1^ts^ (**B**) under SNRK null background. Bottom black bars represent night, white bars represent day. Each row represents a single fly. Within each row, black bar represents duration of immobility. Activation or inhibition was conducted at the second day and majority of flies remained immobile ever since, which were proven to be dead after recording.

## Supplementary Tables

**Table S1. Normalized Sleep Duration in the CCT Screen.**

Normalized sleep duration across 223 Gal4 driver lines in the CCT screen crossed with UAS-Snrk-RNAi during the night, day, and whole day. For each sheet, the first row represents the average sleep duration of the screened driver line; the second row represents the gene name of the screened driver line; the rows below represent the normalized sleep duration of individual flies for the screened driver line. Different driver lines representing the same gene are shown in separate groups. Driver lines are arranged from the lowest to the highest sleep duration.

